# ENDO-Pore: High-throughput linked-end mapping of single DNA cleavage events using nanopore sequencing

**DOI:** 10.1101/2021.07.02.450912

**Authors:** Oscar E. Torres Montaguth, Stephen J. Cross, Kincaid W.A. Ingram, Laura Lee, Fiona M. Diffin, Mark D. Szczelkun

## Abstract

Mapping the precise position of DNA cleavage events plays a key role in determining the mechanism and function of endonucleases. ENDO-Pore is a high-throughput nanopore-based method that allows the time resolved mapping single molecule DNA cleavage events *in vitro*. Following linearisation of a circular DNA substrate by the endonuclease, a resistance cassette is ligated recording the position of the cleavage event. A library of single cleavage events is constructed and subjected to rolling circle amplification to generate concatemers. These are sequenced and used to produce accurate consensus sequences. To identify the cleavage site(s), we developed CSI (Cleavage Site Investigator). CSI recognizes the ends of the cassette ligated into the cleaved substrate and triangulates the position of the dsDNA break. We firstly benchmarked ENDO-Pore using Type II restriction endonucleases. Secondly, we analysed the effect of crRNA length on the cleavage pattern of CRISPR Cas12a. Finally, we mapped the time-resolved DNA cleavage by the Type ISP restriction endonuclease LlaGI that introduces random double-strand breaks into its DNA substrates.

## INTRODUCTION

Endonucleases are diverse group of proteins with a variety of mechanisms to introduce dsDNA breaks. Given their importance for biological processes and their biotechnological applications, endonucleases have been studied extensively. Although catalytic mechanisms for simple dimeric endonucleases are available, there are many complex nucleases that remain poorly understood. Mapping the precise position of DNA cleavage events plays a key role in determining the mechanism and function of this important class of enzymes.

With the recent use of programable nucleases for gene editing, like Cas9 and Cas12a, *in vivo* Next Generation Sequencing (NGS) based methods have been developed for the evaluation of off-target cleavage events (1–3). While these *in vivo* approaches provide valuable information about the specificity of site-directed nucleases, due to cellular repair processes, information about the precise position and nature of the dsDNA break is lost. Traditional *in vitro* gel-based sequencing methods can identify cleavage sites with single nucleotide precision, but their use is limited by low throughput and by being confined to relatively short sections of DNA per lane. Coverage of multiple cleavage locations and characterization of the temporal progression of a cleavage reaction are therefore labour intensive.

Alternative high-throughput next generation sequencing methods have been developed to overcome these limitations (4,5). However, the data obtained remains suboptimal since the ends are sequenced independently, losing information on the link between top and bottom strand reactions at a single cleavage event. Recently, methods based on the use of small circular substrates in combination with paired-end short-read Illumina sequencing have been developed to perform linked-end mapping (6,7). However, the capacity to use small circular substrates can be limited when studying endonucleases that recognize multiple targets over long distances, e.g., the Type I and Type III restriction endonucleases (8–10).

A low throughput linked-end mapping technique based on Sanger sequencing that can utilise plasmid-sized substrates has been described previously and applied to mapping non-specific cleavage by Type I and Type ISP restriction endonucleases (11,12). This approach is based on the ligation of a resistance cassette at the position where the dsDNA break was generated, recording the position of both ends of the cleavage event. Following transformation and antibiotic resistance selection, plasmids can be purified separately from individual colonies and sequenced using either two primers diverging from the resistance cassette or a single primer across the target site. Because each colony is sequenced independently, performing time-resolved mapping of complex nuclease mechanisms can become laborious and expensive.

To improve the method, long read sequencing technologies could be used since they can link the two ends of a dsDNA break in a single read. Oxford Nanopore Technologies sequencing is an easy-to-implement, high-throughput long read sequencing technology that provides flexibility in data acquisition. However, the single read error rate is high, making the base level identification of dsDNA breaks challenging. Different alternatives have been developed to increase single read accuracy (13–15). One of these strategies is the Rolling Circle to Concatemeric Consensus (R2C2) method (13). R2C2 uses circular molecules as a template for rolling circle amplification (RCA) and generates concatemers containing multiple copies of the same sequence. These concatemers are then sequenced, split into individual repeats and consensus sequences with increased base accuracy are generated.

Here we describe ENDO-Pore, a high-throughput nanopore-based method that uses the R2C2 principle to allow the time resolved linked-end mapping of both ends of single molecule DNA cleavage events *in vitro* using plasmid substrates. Firstly, we benchmarked ENDO-Pore by mapping different Type II restriction endonuclease with well described cleavage sites. Secondly, we analysed the effect of crRNA spacer length on the cleavage pattern of *Lachnospiraceae bacterium* Cas12a (LbCas12a). Finally, we mapped the time-resolved DNA cleavage by the *Lactococcus lactis Type ISP restriction endonuclease* LlaGI, an ATP-dependent translocase that can introduce dsDNA breaks into non-specific sequences hundreds of base pairs distant from its recognitions site. ENDO-Pore is an accessible method that is generally applicable to any process that produces a dsDNA break in a circular DNA, and can be used for both cleavage site discovery and mechanistic validation.

## RESULTS

### ENDO-Pore Workflow – Library preparation and nanopore sequencing

ENDO-Pore uses plasmids (or other circular DNA) as substrates for the endonuclease being tested so that the cleaved ends are linked in the linear product. Following DNA linearisation, the DNA ends are repaired and a resistance cassette ligated by TA cloning, recording the position of the cleavage event (Figure 1). If a circular DNA without an origin of replication is to be used, the resistance cassette must include an origin. A library of plasmids representing multiple single cleavage events are produced by transformation, selection of single colonies on antibiotic media and DNA purification from pooled colonies. The number of colonies used determines the number of single cleavage events that will be available for analysis; it is good practice to over-sample the expected number of unique cleavage loci. The library is then subjected to RCA, generating multibranched concatemers (Figure 1). To produce linear fragments suitable for nanopore sequencing, the RCA products are debranched using T7 endonuclease and the concatemers enriched using a size cut-off to favour fragments with multiple repeats (Supplementary Figure S1). The debranched DNA is prepared for sequencing by ligation of DNA barcodes and sequencing adaptors. Sequencing was carried out using Oxford Nanopore Technologies R9.4.1 cells.

**Figure 1.**
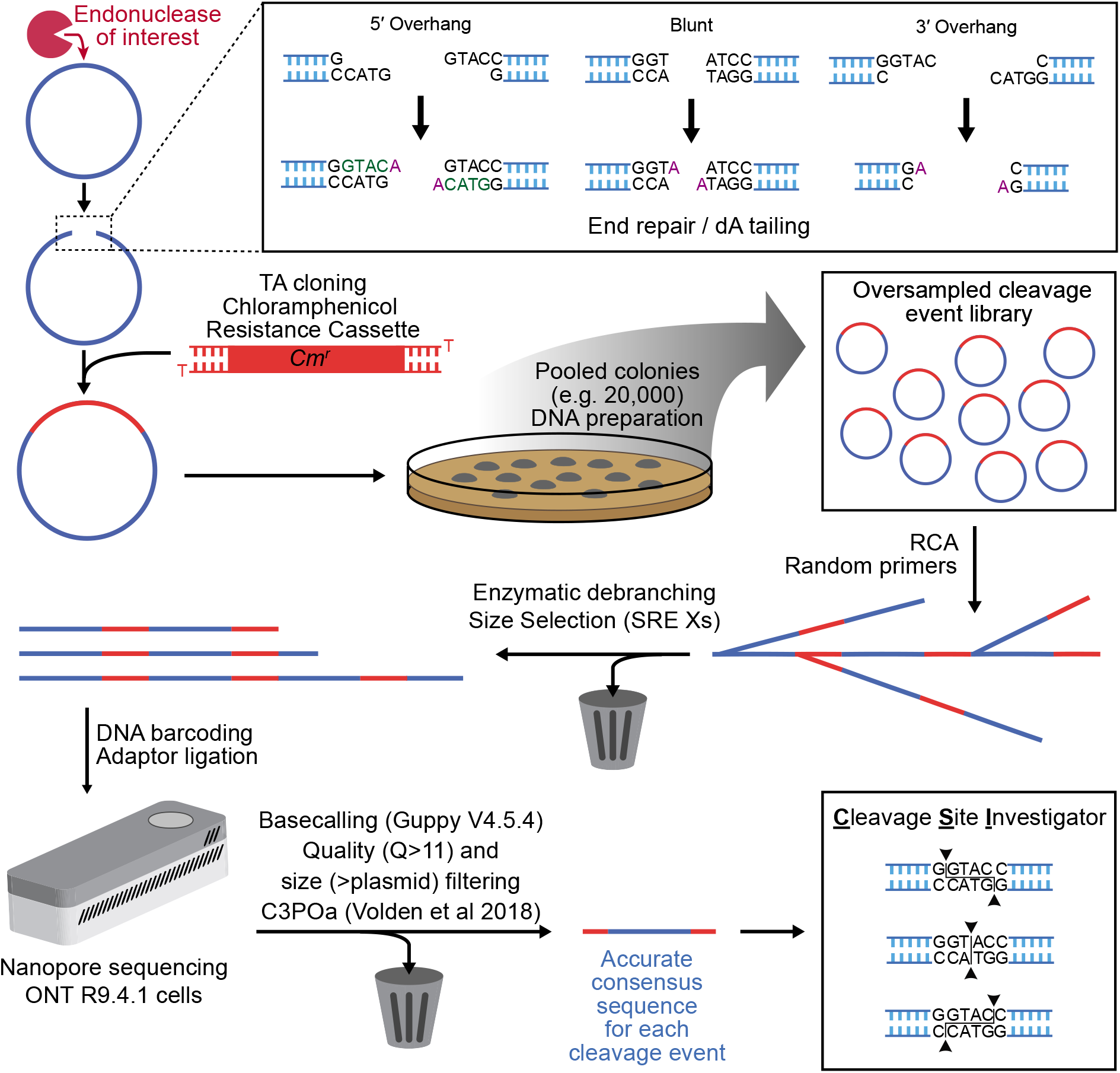
ENDO-Pore Workflow. A circular DNA substrate (blue circle) is digested with the endonuclease of interest (red Pac-Man) and the linear DNA end-repaired (inset). This will fill-in 5’ overhangs and remove 3’ overhangs, while blunt ends remain intact. dA-tailing allows for TA ligation of a resistance cassette (e.g. chloramphenicol used here) that marks the position of the cleavage site. Each individual DNA molecule represents a single cleavage event. Following transformation of *E. coli*, DNA preparation from pooled single colonies produces a DNA cleavage library. Rolling circle amplification is used to both linearise the DNA and to produce concatemer copies. The product is debranched by T7 endonuclease and size selection used to favour DNA with multiple copies of the original plasmid substrates (Supplementary Figure S1). DNA barcodes and sequencing adaptors are ligated, and the DNA sequenced using a MinION. Following basecalling, size selection and consensus generation from the concatemers, the cleavage site for each read is identified by the CSI software (Supplementary Figure S2).

### ENDO-Pore Workflow – consensus generation and cleavage site identification

After barcode indexing and filtering of the data by quality score and length (Materials and Methods), high accuracy concatemeric consensus sequences are generated using C3POa (13) (Figure 1). To then identify the cleavage sites, we developed <u>C</u>leavage <u>S</u>ite <u>I</u>nvestigator (CSI) (Materials and Methods). For each consensus, the 5’ and 3’ ends of the resistance cassette are identified by alignment (default value of 20 nucleotides and 100% sequence identity) (Supplementary Figure S2A). Only sequences containing both ends of the cassette are retained. CSI uses the sequences adjacent to the cassette ends in the consensus (default value of 20 nucleotides) to search the reference sequence (the original plasmid substrate sequence, Supplementary Figure S2B) and pairs with the highest alignment scores are recorded (“aligned sequences”). Because the consensus and reference sequences may not have the same start point, a sequence from the consensus midway between the cassette ends is identified (“mid-flag”, Supplementary Figure S2C). Cleavage positions are then mapped using the aligned sequences (Supplementary Figure S2D): for the 5’ aligned sequence, cleavage is scored at the 3’ end on the opposite strand; for the 3’ aligned sequence, cleavage is scored at the 5’ end on the same strand. The mid-flag is used to orient the cleavage site in the reference so that CSI can correctly identify the type of overhangs produced (5’, 3’ or blunt). After repeating this process with the library, CSI provides the cleavage site frequencies, the dinucleotide frequencies at the nicking site and reports a summary of errors where cleavage sites could not be identified (example in Supplementary Figure S3). Where there are multiple cleavage sites, the data can be output as a matrix table suitable to generate a heatmap or displayed as a strand-linkage plot (e.g., see Figure 4 below).

If the enzyme being tested cleaves each strand just once, then the ENDO-Pore workflow and CSI analysis will return the exact cleavage loci. However, if endonuclease activity produces multiple strand cleavages on a single DNA, then a consequence of the end repair processing (Figure 1) is that CSI will only report the cleavage locus that is closest to the 3’ end of each strand (Figure 2, Supplementary Figure S4). This limitation is true of all sequencing methods that rely on polymerase- and exonuclease-dependent end repair.

**Figure 2.**
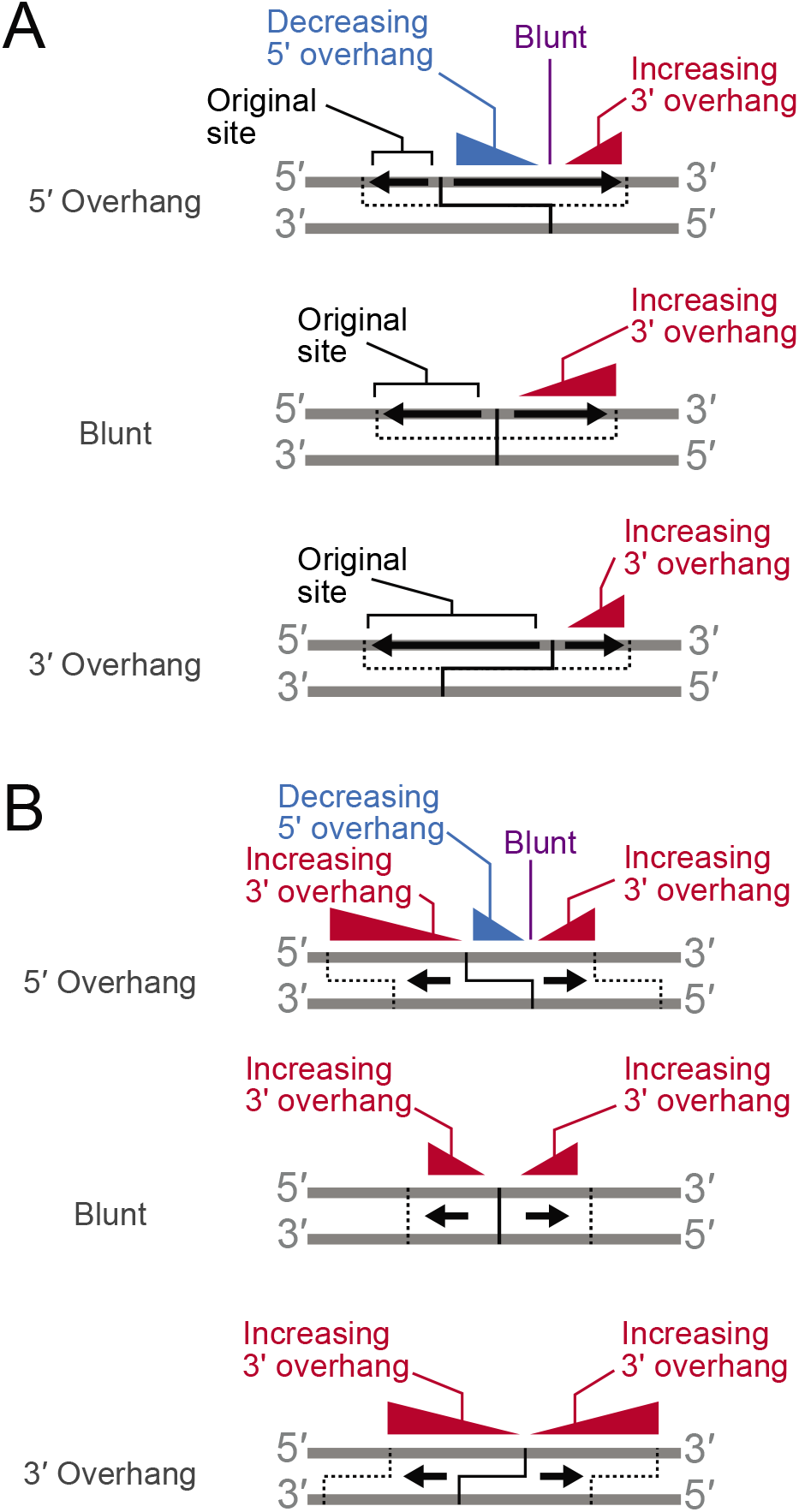
The consequences of DNA end repair are that cleavage sites identified are always those closest to the 3’ end of each strand. (**A**) Consequences for strand-specific processing on the site reported by CSI. (**B**) Consequences for movement and re-cleavage on the site reported by CSI. The solid black lines represent the original dsDNA break and overhangs produced. Black arrows represent movement of the nuclease on one strand (A) or both strands (B). Triangles represent increasing/decreasing apparent spacing between top and bottom stand cleavage loci. Dotted lines represent the cleavage positions and overhangs produced following processing (but not the final cleavage reported). See main text for further details and Supplementary Figure S4 for examples.

Firstly, for enzymes that will cut one or both strands 5’ or 3’ to the initial dsDNA cleavage site, the outcome will depend on the polarity of the secondary cleavage (Figure 2A). For processing in a 5’ to 3’ direction, this will move the cleavage loci towards the 3’ end and tend towards forming 3’ overhangs (Supplementary Figure S4A). If it is suspected that an endonuclease processes the DNA in this way, the temporal progress of the reaction can be followed to deconvolute the cleavage mechanism (12); sampling earlier time point will help to identify the original cleavage site prior to processing. This point is illustrated below in the analysis of the ATP-dependent restriction endonuclease LlaGI. However, for processing in the opposite 3’ to 5’ direction, this will move the cleavage loci towards the 5’ end, resulting in a 5’overhang that will be filled back in during end repair. Hence only the original cleavage site (i.e., closest to the 3’ end) is reported (Figure 2A). This type of processing is therefore invisible to all mapping methods that rely on end repair.

Secondly, the enzyme could move along the DNA (or a second enzyme could bind) and a second dsDNA break will be made at another location with the same spacing between the top and bottom strand cleavages (Figure 2B). Regardless of the identity of the cleavage, this results in an apparent 3’ overhang that can extend over tens to thousands of base pairs. One of the strand cleavage loci could correspond to the original dsDNA break, but this need not be the case (Supplementary Figure S4B). As above, the processing can be deconvoluted by using earlier reaction time points to track the progress of the reaction (12).

### Benchmarking the accuracy of R2C2 method

We first tested the accuracy of the consensus sequences generated by C3POa (13). For this, we used uncut pUC19 as template for RCA and aligned the resulting consensus sequence to the original pUC19 sequence using Minimap2 (16). Since C3POa indicates the number of subreads (“repeats”) used to generate each consensus sequence, we used different repeat cut-offs to test if accuracy increased by excluding concatemers with smaller numbers of subreads. As expected (13), accuracy increased as the repeat cut-off was increased (Figure 3A). The distributions were asymmetric, with a tail of less accurate data that was reduced by increasing the cut-off. Even including all the concatemers produced a distribution with a 25% quartile of 98.2%. A median accuracy of ^~^99.6% with a first quartile of ^~^99.4% could be achieved using a cut-off of ≥5 repeats. Although using higher repeat cut-offs produced increased accuracy, it also produced a reduction in the data output (Figure 3B). As explained below, varying the cut-off can be a useful tool in determining whether an identified cleavage site is a true site or the result of basecalling errors.

**Figure 3.**
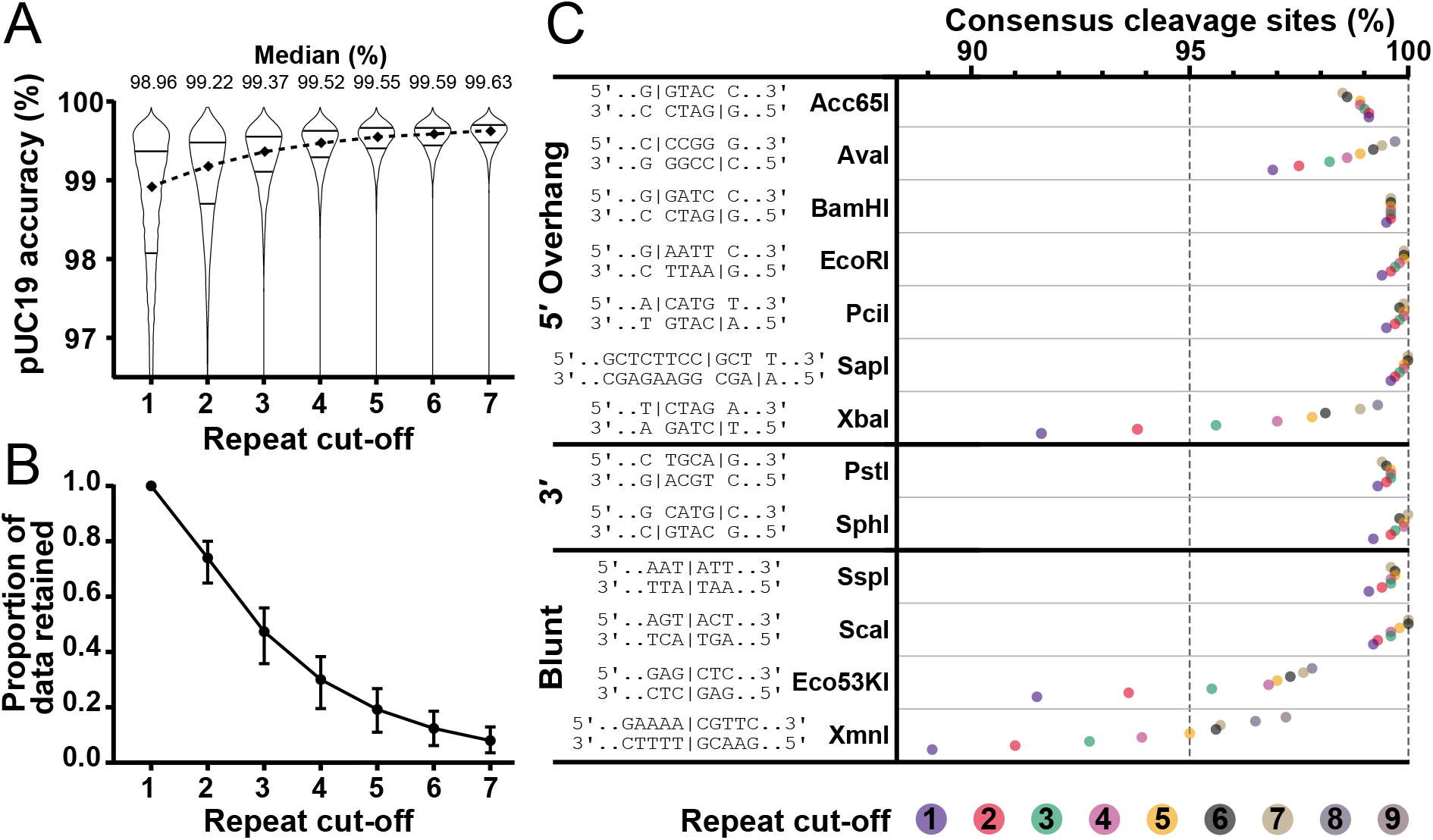
ENDO-Pore Benchmarking. **(A)** pUC19 sequencing accuracy at different repeat cut-offs. Read accuracy was determined by aligning consensus sequences to pUC19 sequence using Minimap2 (16) and is represented by a violin plot for the full dataset at each cut-off. Median accuracy is represented by the diamonds and the quartiles are shown by the horizontal lines. **(B)** Relationship between repeat cut-off and data loss. Average values for the proportion of data retained after different repeat cut-offs were used for the different restriction endonucleases in panel C (points are the average, error bars are the range). **(C)** Validation of ENDO-Pore using commercial Type II restriction endonucleases. Accuracy is represented as the percentage of sequences with the expected consensus cleavage site at different repeat cut-offs, as indicated by the coloured circles. The vertical dotted line at 95% is the “terminal integrity” guaranteed by New England Biolabs determined by ligation and re-cleavage tests.

### Benchmarking the accuracy of ENDO-Pore against commercial Type II restriction endonucleases

To test the accuracy of ENDO-Pore, we tested 13 commercially available Type II restriction endonucleases with established recognition sites that are expected to cut at a single position on each strand (Figure 3C, Supplementary Figure S5) (17). The accuracy of the method is described by percentage of consensus sequences (i.e., events) displaying the expected cleavage site. As the number of subreads used to generate the consensus sequence increased, the percentage of events displaying the expected cleavage site also increased in most cases. This is more evident for sequences with an initial lower accuracy (e.g. AvaI, XbaI, Eco53KI and XbaI). For nine of the thirteen enzymes tested, ENDO-pore exceeded the 95% terminal integrity certified by New England Biolabs, even at the lowest repeat cut-off, and approached 100% as the cut-off was increased. AvaI was less accurate but still exceeded 95% with a repeat cut-off of ≥1 and had 99% accuracy with a cut-off of ≥5. XmnI, EcoK53I and XbaI required elevated repeat cut-offs to exceed 95% and are considered further below.

XmnI accuracy was sub-optimal, with a subread cut-off of ≥5 required to achieve 95% accuracy. The XmnI recognition sequence (5’-GAANN | NNTTC-3’) in the pUC19 substrate used here (Figure 3C) would produce a four adenine repeat at one end of the linear DNA product. After end repair and dA-tailing (Figure 1), an additional adenine would be added, generating a five-nucleotide repeat adjacent to the chloramphenicol cassette after ligation. Homopolymer runs as short as 3 bp can result in apparent INDELs due to errors in nanopore basecalling (13,18). Accordingly, the three most represented non-target cleavage events identified by CSI were deletions of 1, 2 or 3 base pairs in the five-adenine repeat (Supplementary Figure S6). These errors were iteratively reduced by increasing the subread cut-off. This is characteristic of sequencing errors, whereas real cleavage events or errors in end-processing or library would increase in frequency. We note that basecalling-dependent insertion errors would produce a longer poly-dA repeat that would fail the CSI search, and hence we did not observe them. The slightly reduced accuracy with AvaI also most likely reflects the homopolymeric dC and dG runs in the sequence which were again reduced iteratively by increasing the sub-read cut-off.

For EcoK53I and XbaI, the highest frequency non-target sites identified by CSI were consistent with loss of an AG dinucleotide from one or other side of the site (Supplementary Figure S6). Following dA-tailing and ligation to the chloramphenicol cassette, the sequence at the ligation point would be 5’-AGAGA-3’. It appears that this repeated sequence is miscalled with loss of an AG dinucleotide. Since this error was systematically reduced by increasing the sub-repeat cut-off (Figures 3C and Supplementary Figure S6), the error is likely a sequencing error rather than relaxed endonuclease cleavage or misprocessing during library production.

Because of over-sampling of the Type II cleavage events, we also observed additional sequences with low frequency (<0.5%) that could not be readily explained as basecalling errors. Some of these were due to indexing errors that are a consequence of misidentifying barcode sequences or sample contamination (e.g., a ScaI cleavage event identified in the EcoK53I data, Supplementary Figure S6). Other events represented cleavage at the consensus site and at a second random site. These could be due to either a low background of linear DNA in the plasmid preparation that was cut by the restriction enzyme, or relaxed cleavage at a second off-target site (“star activity”) (19). These events were often spaced hundreds of base pairs apart, leading to large deletions with apparent 3’ overhangs as called by CSI. Since these events shorten the plasmid, they are over-represented in longer concatemers selected by an increase in the repeat cut-off. The reduction in accuracy seen with Acc65I, PstI and SspI with repeat cut-offs of ≥5 and above could be explained by an over-representation of deletion events.

In summary, based on the restriction endonuclease benchmarking, ENDO-Pore provides a reliable method to map DNA cleavage sites. By increasing the repeat cut-off, accuracy of >99.5% can be achieved and where there are problems due to sequencing errors, these can be identified. However, it should be noted that increasing the cut-off also reduces the amount of data, and low N values can lead to misinterpretation of events.

### Benchmarking ENDO-Pore against CRISPR-Cas12a and crRNA with varying spacer length

To explore the ability of ENDO-Pore to map variation in cleavage loci at a specific sequence due to enzyme mechanism, we analysed the effect of crRNA spacer length on the cleavage pattern of *Lachnospiraceae* bacterium ND2006 Cas12a (LbCas12a). LbCas12a requires a single CRISPR RNA (crRNA) to introduce double strand breaks into its target DNA. crRNAs contain a spacer sequence complementary to the target site that is the responsible for the specificity of the enzyme (20,21) (Figure 4A). First, a Cas12a-crRNA complex binds DNA through interactions with the T-rich Protospacer Adjacent Motif (PAM) (22). This initial interaction leads to the ATP-independent unwinding of the target DNA and the formation of an R-loop in which a heteroduplex is formed between the target strand (TS) and the crRNA spacer sequence (23,24). R-loop formation allows positioning of the non-targeted strand (NTS) in the RuvC active site and NTS nicking. The cleaved ends are released but can re-enter the active site multiple times, where new cleavage events lead to gap formation (25). Following NTS processing, conformational transitions lead to the TS entering the active site, with cleavage producing a 5’ staggered dsDNA break.

**Figure 4.**
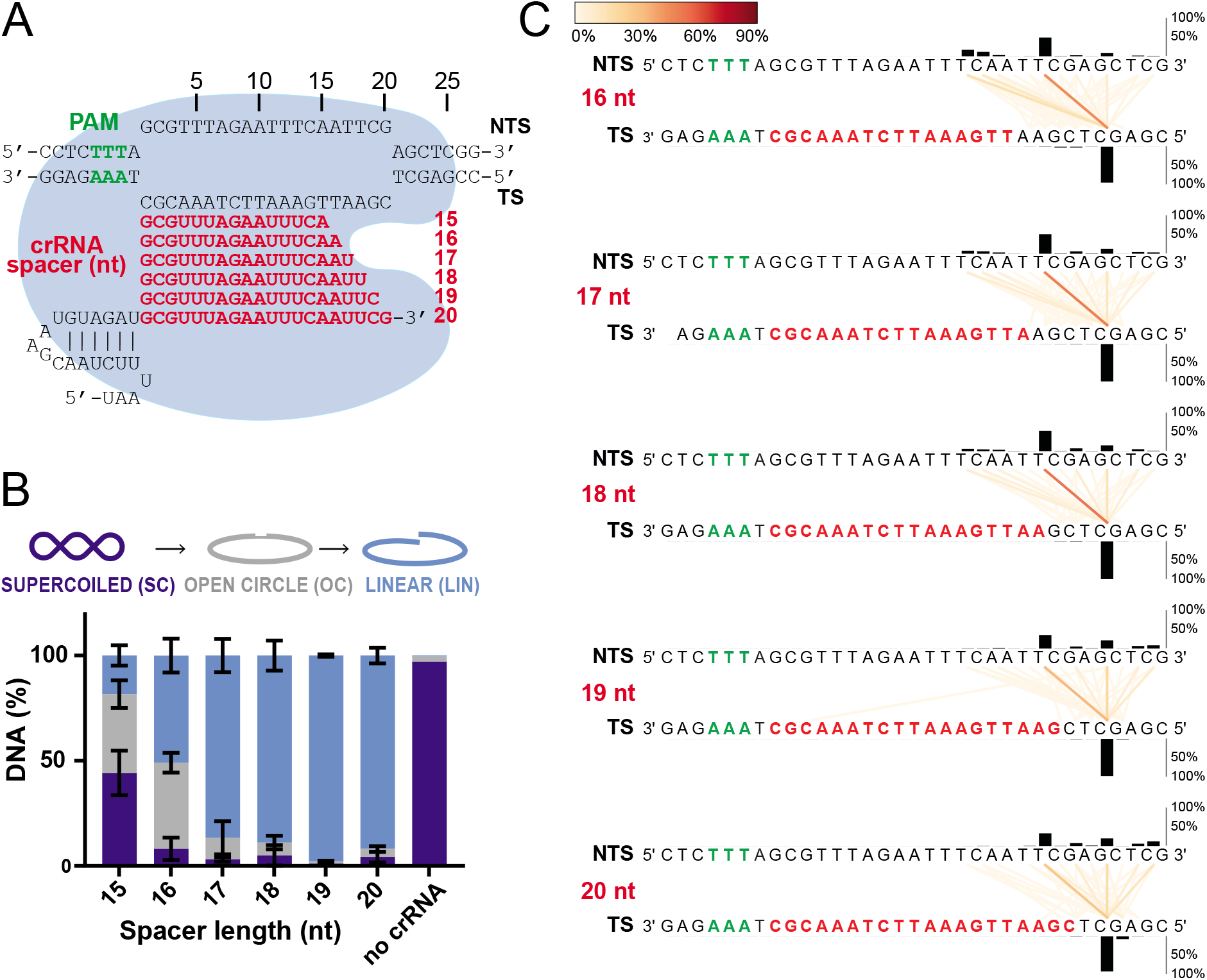
The effect of crRNA spacer length on the cleavage pattern of CRISPR-Cas12a. (**A**) R-loop formation between the plasmid target and Cas12a crRNA with 15-20 nt spacer sequences (red). The PAM is shown in green and the bi-lobed Cas12a structure represented by the blue shape. (**B**) Plasmid DNA cleavage after 2 hours. Supercoiled (SC) substrate, nicked (open circle, OC) intermediate and linear (LIN) product were separated by agarose gel electrophoresis (Supplementary Figure S7) and the tritium-labelled DNA quantified by scintillation counting. Errors bars are SD from 3 repeat experiments. (**C**) Strand-linkage plots (SLPs) generated by CSI. Each SLP displays the relative frequency of cleavage events (represented as a connecting line between the NTS and TS according to the heatmap) and the frequency of strand-specific cleavage events above or below the relevant strand (black histograms).

It has been noted previously that Cas12a crRNA with spacer sequences of 15-16 nucleotides (nt) can still support DNA cleavage, although the sites of cleavage can vary (4,26). To analyse this effect using ENDO-Pore, we initially tested DNA cleavage using 6 different crRNA with spacers ranging from 15 – 20 nt truncated at the PAM distal 3’ end (Figure 4A). Since less than 20% of the DNA was linearised using the 15 nt crRNA after 2 hours (Figure 4B, Supplementary Figure S7), we did not analyse this ribonucleoprotein (RNP) further. For the other crRNA, DNA was prepared for ENDO-Pore sequencing. The data was analysed with a repeat cut-off of ≥5; increasing the repeat cut-off did not vary significantly the results observed. The data is presented as strand-linkage plots (SLPs) generated by CSI that link the cleavage loci of the top and bottom strands with vertical/diagonal lines. The frequency of dsDNA cleavage events is represented by optional width and colouring of the lines according to a heat map, and for the strand-specific cleavage events by the bar graphs above or below the relevant strand (Figure 4C).

Regardless of the spacer length, the position of the TS nicking event was unvarying, with more than 90% of events taking place at position 22, at the PAM distal end beyond the end of the protospacer (Figure 4C). In contrast, although all crRNA produced a principal product at position 18 on the NTS, the locations of other cleavage events were more variable, generating apparent ends with different overhangs. Previous mapping experiments with Cas12a (4,26) have suggested that crRNA with shorter spacers (16 – 18 nt) promote NTS cleavage events closer to the PAM, resulting in elongated 5’ overhangs of 7-6 nucleotides. We also observed elevated frequencies of cleavage events at positions 13, 14 and 15 for crRNA with 16-18 nt spacers (corresponding to 5’ overhangs of varying length), and these events became less frequent as the spacer was lengthened to the full R-loop (20 nt) and were replaced by increased cleavage at positions 20, 22, 24 and 25. As we note (Figure 2), blunt ends and 3’ overhangs can result from processing of initial 5’ overhangs, which is likely the case here since gap formation is key to the handover between the NTS and TS in the sequential cleavage pathway (25). Our data would suggest that gap formation in the 5’ – 3’ direction following the initial NTS cleavage is support by crRNA with longer spacer sequences, which is presumably a consequence of a longer R-loop.

### Benchmarking ENDO-Pore against a Type ISP restriction endonuclease with variable cleavage over hundreds of base pairs

Finally, to test the performance of ENDO-Pore with an endonuclease system that produces a pattern of cleavage events distributed over longer DNA distances, we analysed the *Lactococcus lactis Type I single polypeptide (SP) restriction endonuclease* LlaGI (8,12,27). LlaGI is a single polypeptide, multidomain protein with modification and restriction activities undertaken by the same monomeric protein. It has four domains: A DNA target recognition domain (TRD) that recognizes the sequence 5’-CTnGAyG-3’ (where n is A, G, C or T and y is C or T), an *S*-adenosyl methionine-dependent methyltransferase domain, an SF2-helicase-like ATPase motor domain and an mrr-like nuclease domain (27). Once the TRD recognizes its target site, a conformational change promotes unidirectional, ATP-hydrolysis-dependent translocation downstream from the site. On DNA with a pair of sites in inverted head-to-head repeat, the collision of two converging enzymes leads to nicking of the top strand by one nuclease domain and of the bottom strand by the other nuclease domain. Movement of the collision complex and continued strand-specific DNA nicking eventually results in the formation of dsDNA breaks (12) (Figure 5A).

**Figure 5.**
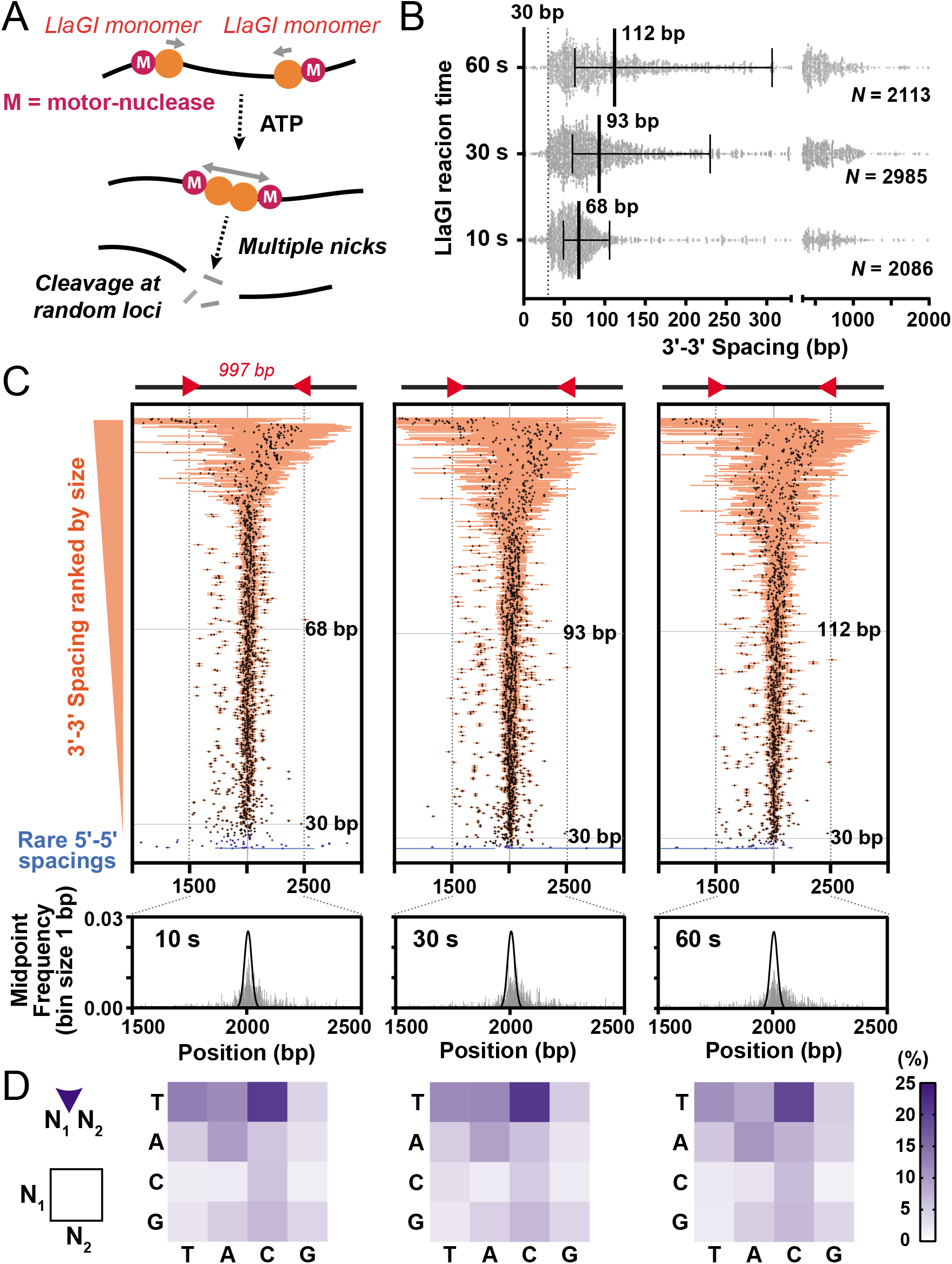
Mapping the long-range communication and cleavage by the Type ISP restriction endonuclease LlaGI. (**A**) Cleavage mechanism of LlaGI. (**B**) Scatter plot of the 3’-3’ spacing between nicking events reported by CSI. Error bars represent the interquartile distance; thick bars represent the median values; N represents the total number of cleavage events analysed per time point. **C.** Tornado plots displaying individual events ranked by spacing between nicking events. Horizonal orange (3’ overhangs) or blue (5’ overhangs) bars represent the spacing between the nicks closest to the 3’ end of each strand. Crosses indicate the midpoint of each event. Positions of the 30 bp and median spacings are shown by grey horizontal lines. (*lower panels*) The frequency of the cleavage midpoints for each time point are represented as grey vertical bars (bin size 1 bp). The black solid lines represent the theoretical collision distribution at time → ∞ of two motors that initiated at a pair of sites a distance *d* = 997 steps apart, given by *d*!/(*n*!·(*d* – *n*)!·2^*d*^), where *n* is the position between the sites (28). **D.** Dinucleotide cleavage frequency for individual time points calculated by CSI.

The LlaGI DNA cleavage reaction was previously mapped using the low throughput Sanger-sequencing mapping method of Bickle and coworkers (11) using a DNA with a relatively short inter-site spacing of 97 bp and limited to ^~^100 events per time point (12). Here we used the high-throughput advantage of ENDO-Pore to map the time-resolved cleavage of a plasmid substrate in which the two head-to-head target sites were 997 bp apart. For this test, we took samples of a cleavage reaction after 10, 30 and 60 s (Supplementary Figure S8). The distributed cleavage products identified by CSI can be visualised in several ways. Firstly, an SLP emphasises that most cleavage events produced 3’ overhangs, even at the earliest time points, and that cleavage occurs both between the sites and up- and down-stream of the sites, across almost the entire length of the plasmid (Supplementary Figure S9). The main 3’ - 5’ overhang distances were widely distributed, with a median that increased as time progressed (Figure 5B) and showing distinct minima at 30 bp. To map the midpoints of the cleavage events (that can represent the initial points of motor collision for symmetrically processed sites), we ranked the data by overhang length and plotted the midpoint of collision on the DNA with the overhang length as error bars (the “tornado” plots in Figure 5C). The distribution of cleavage midpoints could then be calculated (Figure 5C, lower panels). Finally, given that the cleavage occurs away from the recognition site at non-specific loci, the dependence of cleavage on local sequence could be judged from the dinucleotide frequencies calculated by CSI (Figure 5D).

In the collision-cleavage model for LlaGI, the initial complex produced upon collision between converging motors places the nuclease domains at too great a distance for strand-specific nicks to produce a dsDNA break (9,12). Subsequent up and downstream motion of the complex and continued nicking leads eventually to two closely spaced nicks and a dsDNA break. The spacing data collected here (Figure 5B) is consistent with this model and provides more statistical certainty than the previous low-throughput data. The 30 bp minimum is again observed. Note that this does not represent the closest spacing of nick sites since CSI will report the cleavage loci nearest to the 3’ end on both strands (Figure 2). Instead, it represents the minimum distance for the outermost nicks produced at the initial collision; the dsDNA breaks most likely resulted from additional nicking between these points. As time progresses, continued nicking leads to an increasing distance between the outermost nicks (Figure 5B).

The widening of the outermost nicks with time is also observed in the tornado plots as a time-dependent increase in error bar width (Figure 5C). These plots also show that most collisions occur approximately midway between the sites. This was not observed previously with the 97 bp spacing (12), but is expected from mapping of cleavage products on longer spacings using agarose gels (8) and from theoretical expectations of the collision of two converging stepping motors (28). The distributions of the midpoints do not change markedly with time (Figure 5C, lower panels). This is consistent with the model that following collision the complex moves both up- and downstream with equal probability, while nicking the DNA. Pushing backwards of one motor by the other would result in a shift in the midpoint. Some of these events appear in the longer spacings that are also observed at the earliest time points. There is some asymmetry in the distribution of these longer spacings; it is not clear as to the source of this asymmetry, although it should be noted that the plasmid origin overlaps with some potential cleavage loci and cleavage in this region will not show up in the mapping data (Supplementary Figure S9).

For some shorter cleavage spacings, collisions occurred away from the midpoint (Figure 5C, Supplementary Figure S9). These were most likely due to dissociation of one converging motor during translocation. Since other motors can initiate at the recognition site, collisions will still occur but at off-centre positions (28). There are also some collision events that map to the sites, where collision occurred before one of the motors could initiate, and one or two events that are upstream of a site, suggesting that a collision occurred at the site that immediately displaced the site-bound enzyme upstream before motion stalled and cleavage was activated.

An analysis of the frequency of the nucleotides at the nicking positions indicates that the nuclease domain has a dinucleotide preference that does not change appreciably with time (Figure 5D). A similar distribution was observed previously (12), but because of the low throughput of the Sanger method, this earlier data was averaged across three time points.

## DISCUSSION

Here we describe the ENDO-Pore sequencing method that allows high accuracy mapping of DNA cleavage events *in vitro*. The method can be used to map specific, semi-specific and non-specific endonucleases. By using circular DNA substrates, connection can be maintained between both ends of each individual dsDNA break. Such linked-end mapping provides important information that might otherwise be lost, especially when the nicking events on the two strands occur at variable positions. Long read nanopore sequencing allows the use of plasmid-length substrates, providing advantages in terms of ease of production, sequence diversity, and the ability to vary supercoiling that can affect endonuclease cleavage activity (29). Since they are commonly used as substrates for other biochemical characterization methods, ENDO-Pore results can be easily integrated with data generated by different approaches providing a more comprehensive mechanistic overview. Although we benchmarked the method using enzymes with known recognition sites, it would be straightforward to modify the method to search for recognition sites with an uncharacterised enzyme, e.g., by using a plasmid library as templates (e.g., (30)).

Nanopore technology is more accessible and cheaper than other deep sequencing methods, and allows simple user control of data acquisition, e.g., fewer sequences can be collected for a simple Type II enzyme compared to a more complex Type ISP enzyme. Although the single read base accuracy of nanopore sequencing can be a limiting factor, we found that this was significatively improved using the R2C2 method (13). However, as previously described, nanopore sequencing is more error prone with certain sequences, such as homopolymers in which deletions of single repeats are the most common error observed (13,18). Therefore, discerning between potential sequencing errors and real cleavage events remains important for result interpretation. Given that increasing the repeat cut-off improves the accuracy at problematic sequences, this analysis can be performed systematically to look for changes of event frequency distribution. New developments in nanopore sequencing can provide an increase in accuracy at the single read level, e.g., flow cells with improved pores that increase homopolymer accuracy (ONT R10.3), new chemistries that improve translocation rates across the pores, and new robust basecalling algorithms. Additionally, improvements can be made in the quality of DNA preparations (e.g., excluding any linear DNA) and in indexing of barcodes (e.g.,(31)) to reduce spurious results.

Similarly to other next generation sequencing methods for dsDNA break mapping that use end repair as part of the workflow, ENDO-Pore reports on the cleavage loci closest to the 3’ end on each strand (Figure 2). For enzymes with end processing after initial cleavage, end repair can thus result in loss of information about the primary cleavage event. This implies that while 5’ overhangs and blunt ends can be determined with higher confidence, 3’ overhangs might arise from sequential cleavage events and/or end processing. Performing time courses and/or using physical conditions to slow the reaction (e.g., lower reaction temperatures) can help to distinguish between primary and secondary events. For example, by comparing the temporal reactions of LlaGI, linked-end mapping allowed the identification of initial cleavage events and subsequent end processing during a complex reaction over hundreds of base pairs. Nonetheless, 3’ - 5’ processing of nick sites can never be observed (as the DNA is repaired to the original cleavage site) (Figure 2). Therefore lower-throughput gel-based methods can still provide value when used side-by-side with ENDO-Pore to explore the full range of end processing events.

## MATERIALS & METHODS

### Restriction enzyme cleavage reactions

pUC19 was digested using thirteen commercially available Type II restriction enzymes (New England Biolabs). For BamHI, EcoRI, SspI, ScaI and SphI, High-Fidelity (HF) versions were used. Reactions were set up using the restriction buffer and enzyme quantities suggested by New England Biolabs. Reactions were incubated for 1 hour at 37 °C and DNA was purified using the DNA Clean & Concentrator-25 kit (Zymo Research) as described by the manufacturer.

### LbCas12a cleavage reactions

LbCas12a was purified as previously described (29). crRNAs were synthesised and HPLC-purified by IDT (Supplementary Figure S10). For ribonucleoprotein (RNP) complex assembly, 250nM Cas12a and 250nM crRNA were mixed in buffer RB (10 mM Tris-Cl, pH 7.5, 100 mM NaCl, 10 mM MgCl_2_, 0.1 mM DTT, 5 μg/ml BSA) supplemented with 0.05 U/μL SUPERase-In RNase Inhibitor (ThermoFisher) and incubated at 37 °C for 1 hour. 5 nM pSP1 (32) was pre-heated in Buffer RB at 37 °C for 5 minutes. Reactions were started by addition of 50 nM Cas12a RNP and incubated for 2 hours at 37 °C. The reaction was quenched by adding binding buffer from the DNA Clean & Concentrator-25 kit (Zymo Research) supplemented with 30 mM EDTA and 37 mM sodium acetate, and the DNA purified as described by the manufacturer.

### LlaGI cleavage reactions

LlaGI was purified as previously described (27). A modified version of pRMA03 (27) was used as a substrate. pRMA03 was digested using SmaI and PshAI and re-ligated generating pRMA03S, a substrate in which the two LlaGI sites are in head-to-head orientation separated by 997 bp. A reaction mix containing 2 nM pRMA03S, 200 nM LlaGI in TMD Buffer (50 mM Tris-Cl, pH 8.0, 10 mM MgCl_2_, 1 mM DTT) was pre-incubated at 25 °C for 5 minutes. Cleavage reactions were initiated by the addition of 4 mM ATP, and stopped at 10, 30 or 60 s by adding binding buffer from the DNA Clean & Concentrator-25 kit (Zymo Research) supplemented with 30 mM EDTA and 37 mM sodium acetate. Purification of cleaved DNA proceeded as described by the manufacturer.

### Preparation of the chloramphenicol resistance cassette

A chloramphenicol resistance cassette flanked by Hpy188I restriction sites was amplified by overhanging-end PCR from the pACYC184 plasmid (33) using oligos 5’-CTAGCT<u>TCAGA</u>AGTCAGCGACTCGCATCACGCACCAATAACTGCCTTAAAAAAATTACGC-3’ and 5’-TACGTA<u>TCAGA</u>GACGTAGCGTACGCATCGTCAGCGAAAATGAGACGTTGATCGGC-3’. Template plasmid was eliminated by treating with DpnI for 1 hour at 37 °C. The PCR product was purified using DNA Clean & Concentrator-25 kit (Zymo Research). Purified PCR product was digested with Hpy188I (New England Biolabs) for 2 hours at 37 °C to generate single dT 3’ overhangs, and purified.

### ENDO-Pore workflow 1 – library preparation and DNA sequencing

A step-by-step protocol is included in the Supplementary Information. In brief, 1 μg of purified DNA from a cleavage reaction was end-repaired and dA-tailed using the NEBNext^®^ Ultra™ II End Repair/dA-Tailing Module (New England Biolabs). The DNA was purified and ligated with the dT-tailed chloramphenicol cassette. OmniMAX™ 2 T1R *E. coli* cells (ThermoFisher) were transformed with the ligation reactions and single colonies selected using 34 μg/ml chloramphenicol on LB-agar. Cleavage event libraries were generated by pooling colonies followed by plasmid purification. Rolling circle amplification (RCA) was then performed using 10 ng of the cleavage library as a template with EquiPhi29™ DNA Polymerase and exonuclease resistant random hexamers (ThermoFisher). Reactions were incubated for 2 hours at 45 °C, heat inactivated for 10 minutes at 65 °C, and the DNA purified using AMPure XP beads (Beckman Coulter). RCA products were debranched using 10 units T7 Endonuclease/μg DNA (New England Biolabs) for 15 minutes at 37 °C. The debranching reaction was stopped by incubating with 0.8 units Proteinase K (New England Biolabs) for 5 minutes at 37 °C. Debranched products were purified using AMPure XP beads followed by size selection using the Short Read Elimination XS kit (Circulomics). Samples were prepared for Nanopore sequencing using the Ligation Sequencing Kit SQK-LSK109 combined with the Native Barcoding Expansion kits EXP-NBD104/114 (Oxford Nanopore). DNA was sequenced using a MinION and R9.4.1 cells (Oxford Nanopore).

### ENDO-Pore workflow 2 – data processing and Cleavage Site Investigator

Raw reads were basecalled and demultiplexed using Guppy V4.5.4. Reads were filtered for quality (Q > 11) and size (>2000 bp) using NanoFilt (34). Circular concatemeric sequences were generated using C3POa v2.2.2 (13). Individual dsDNA breaks were identified using Cleavage Site Investigator (CSI). The principle of the method is described in the main text and in Supplementary Figure S2. The code for CSI, a manual and test data are available at https://doi.org/10.5281/zenodo.5057043. The manual is also included in the Supplementary Information.

## Supporting information

Manual for the CSI software

Wet lab protocol

## DATA AVAILABILITY

The code for CSI, a manual and test data are available at https://doi.org/10.5281/zenodo.5057043. The consensus, cassette and reference sequences used in this study are available at https://doi.org/10.5523/bris.367vrebu1ee2a23ro8gy6ggfpv.

## ACKNOWLEDGEMENTS

We thank Tom Gorochowski and James Graham for nanopore advice and help with materials, and Roger Volden for help with C3POa.

