## Supplementary material for "ENDO-Pore: High-throughput linked-end mapping of single DNA cleavage events using nanopore sequencing": Manual for the CSI software

### CSI: Cleavage Site Investigator

### Features

- Run straight from command line
- Compatible with FASTA file format (.fa and .fasta)
- Determine top and bottom strand cleavage events
- Export results to .csv files
- Create visual event distributions as heatmaps and strand linkage plots

### Contents

- Features
- Contents
- Installation
- Usage
  - Notes
  - Running CSI
  - Generating strand linkage plots (SVG)
  - Generating heatmap plots (CSV)
  - Generating heatmap plots (SVG)
- Output file formats
  - CSI summary file
  - CSI individual results file

### Installation

1. Download CSI source code from GitHub
2. Install Python (tested with Python 3.9.1)
3. Install required libraries (BioPython, Seaborn, SVGWrite and TQDM)
   - Either using Pip

```
pip install biopython==1.79
pip install tqdm==4.55.1
pip install seaborn==0.11.1
pip install svgwrite==1.4
```

   - Or using the provided Anaconda environment file ("csi.yml" in "resources" folder)

```
conda env create -f csi.yml
```

### Usage

#### Notes

- Example files for testing CSI are included in the "data" folder of this repository. The files are from the full data set found here. These files are:
  - "ex_cassette.fa" - Cassette sequence (must contain one sequence). Example file is for "Splint1TA".
  - "ex_consensus.fa" - Consensus sequence(s) (can contain multiple sequences). Example file is a subset of sequences from "Cas12a_17.fa" sample.
  - "ex_reference.fa" - Reference sequence (must contain one sequence). Example file is for "CrisprplasR".
- The above files are used throughout the following code demos.
- Each program (csi.py, heatmapcsv.py, heatmapsvg.py and strandlinkageplot.py) can be run entirely from command line. Full argument documentation is accessible using the `-h` (or `--help`) flag (e.g. `python csi.py -h`).

#### Running CSI

##### Basic use

- The main CSI program is run using csi.py. This will analyse the specified consensus sequences and optionally output event distributions, summary statistics and plots (advanced plotting options available by running heatmapsvg.py and strandlinkageplot.py directly).
- CSI requires a minimum of three arguments, specifying paths to the cassette (`-ca` or `--cassette_path`), reference (`-r` or `--reference_path`) and consensus (`-co` or `--consensus_path`) files.
- The following command is an example

```
python .\src\csi.py -ca .\data\ex_cassette.fa -r .\data\ex_reference.fa -co .\data\ex_consensus.fa
```

- With default parameters (no optional arguments specified) a basic summary will be displayed with the following sections:

| Label | Description |
| --- | --- |
| "TS position" | Position of the top-strand cleavage event |
| "BS position" | Position of the bottom-strand cleavage event |
| "Split seq" | `True` if the cleavage event spanned the start/end of the reference sequence, `False` otherwise |
| "Count" | Number of identified events matching this cleavage event (% of total identified events shown in parenthesis) |

| Label | Description |
| --- | --- |
| "Type" | Type of cleavage event (either "Blunt end", "3′ overhang" or "5′ overhang") |

- An example output is shown below:

```
RESULTS:
    Full sequence frequency:

        TS position: 1289
        BS position: 1293
        Split seq:   False
        Count:       396/787 (50.3% of events)
        Type:        5' overhang

        TS position: 1293
        BS position: 1293
        Split seq:   False
        Count:       97/787 (12.3% of events)
        Type:        Blunt end

        TS position: 1284
        BS position: 1293
        Split seq:   False
        Count:       67/787 (8.5% of events)
        Type:        5' overhang

        ...
```

##### Advanced control

- CSI offers optional command line parameters to specify execution settings (e.g. the number of bases to fit) as well as additional outputs (e.g. summary CSV files or rendered heatmap plots).

| Argument | Description | Default value |
| --- | --- | --- |
| `-h`, `--help` | Show help message (lists all required and optional arguments). | NA |
| `-rf`, `--repeat_filter` | Expression defining filter for accepted number of repeats. Uses standard Python math notation, where 'x' is the number of repeats (e.g. 'x>=3' will process all sequences with at least 3 repeats). | NA |
| `-lr`, `--local_r` | When grouping sequences at restriction sites, this is the half width of the local sequences to be extracted. For example, for a sequence 5′...AAT\|ATT...3′, `-lr 1` would yield "TA", whereas `-lr 2` would yield "ATAT". | 1 |

| Argument | Description | Default value |
| --- | --- | --- |
| `-mg`, `--max_gap` | Maximum number of nucleotides between 3′ and 5′ restriction sites. | 10000 |
| `-mq`, `--min_quality` | Minimum match quality. Specified in the range 0-1, where 1 is a perfect match. | 1.0 |
| `-nb`, `--num_bases` | Number of bases to match when comparing sequences (e.g. when searching for cassette ends in a consensus sequence). | 20 |
| `-pr`, `--print_results` | Prints results to the terminal once a complete file has been processed. | NA |
| `-en`, `--extra_nt` | Number of additional nucleotides to be displayed either side of the cleavage site (when `-pr` or `--print_results` is specified). | 0 |
| `-sp`, `--show_plots` | Display plots showing local sequence distributions as a heatmap and pie-chart. | NA |
| `-wslp`, `--write_strandlinkageplot` | Write strand linkage plot image to SVG file. Output file will be stored in consensus file folder with same name as the consensus file, but with the suffix '_strandlinkageplot'. To generate strand linkage plots with greater control over rendering, see Generating strand linkage plots (SVG) | NA |
| `-whsa`, `--write_heatmap_svg_auto` | Write heatmap image (only spanning range of identified event positions) to SVG file. Output file will be stored in consensus file folder with same name as the consensus file, but with the suffix '_heatmap'. To generate heatmaps with greater control over rendering, see Generating heatmap plots (SVG). | NA |
| `-whsf`, `--write_heatmap_svg_full` | Write heatmap image (spanning full range of reference sequence) to SVG file. Output file will be stored in consensus file folder with same name as the consensus file, but with the suffix '_heatmap'. To generate heatmaps with greater control over rendering, see Generating heatmap plots (SVG). | NA |
| `-whca`, `--write_heatmap_csv_auto` | Write heatmap image (only spanning range of identified event positions) to CSV file. Output file will be stored in consensus file folder with same name as the consensus file, but with the suffix '_heatmap'. To generate heatmaps with greater control over rendering, see Generating heatmap plots (CSV). | NA |
| `-whcf`, `--write_heatmap_csv_full` | Write heatmap image (spanning full range of reference sequence) to CSV file. Output file will be stored in consensus file folder with same name as the consensus file, but with the suffix '_heatmap'. To generate heatmaps with greater control over rendering, see Generating heatmap plots (CSV). | NA |

| Argument | Description | Default value |
| --- | --- | --- |
| `-wi`,`--write_individual` | Write individual cleavage results to CSV file. Output file will be stored in consensus file folder with same name as the consensus file, but with the suffix '_individual'. For more information on the individual results file format, see CSI individual results file. | NA |
| `-ws`, `--write_summary` | Write summary of results to CSV file. Output file will be stored in consensus file folder with same name as the consensus file, but with the suffix '_summary'. For more information on the summary results file format, see CSI summary file. | NA |
| `-wo`, `--write_output` | Write all content displayed in console to a text file. Output file will be stored in consensus file folder with same name as the consensus file, but with the suffix '_output'. | NA |
| `-ad`, `--append_datetime` | Append time and date to all output filenames (prevents accidental file overwriting). | NA |
| `-v`, `--verbose` | Display detailed messages during execution. | NA |

#### Generating strand linkage plots (SVG)

##### Basic use

- Strand linkage plots can be exported to SVG directly from CSI summary and individual results files using strandlinkageplot.py.
- At a minimum, strandlinkageplot.py requires arguments specifying the path to a CSI summary or individual results file (`-d` or `--data_file` argument) and the output SVG path (`-o` or `--out_path` argument).
- For example, the following command will generate a strand linking plot using default parameters:

```
python .\src\strandlinkageplot.py -d .\data\ex_consensus_summary.csv -o
.\data\output_strandlinkageplot.svg
```

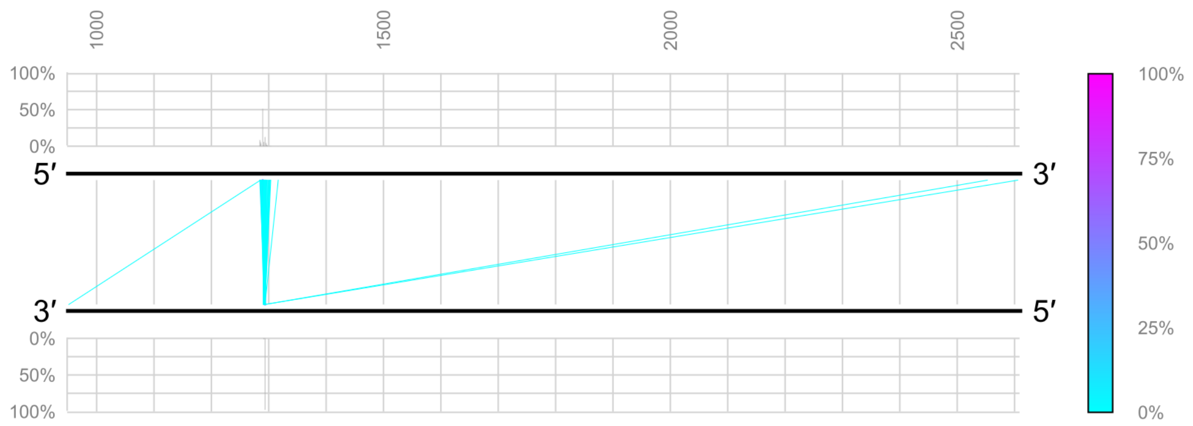


#### Advanced control

- To afford greater control over various aspects of plot rendering, strandlinkageplot.py accepts over 50 different command line arguments. Full descriptions for these arguments can be viewed using the `-h` or `--help` flag.
- The following figure uses optional arguments to zoom in on a specific sequence region (`-pr 1260 1320`), applies closer grid spacings (`-g_i 10 -gl_i 10`), uses a different colourmap (`-e_c plasma`) and displays the DNA as a letter sequence (`-d_m seq`; Note: this requires the reference sequence to be provided via `-r`):

```
python .\src\strandlinkageplot.py -d .\data\ex_consensus_summary.csv -o
.\data\output_modified_plot.svg -e_c plasma -pr 1260 1320 -g_i 10 -gl_i 10 -d_m
seq -r .\data\ex_reference.fa -d_s 12
```

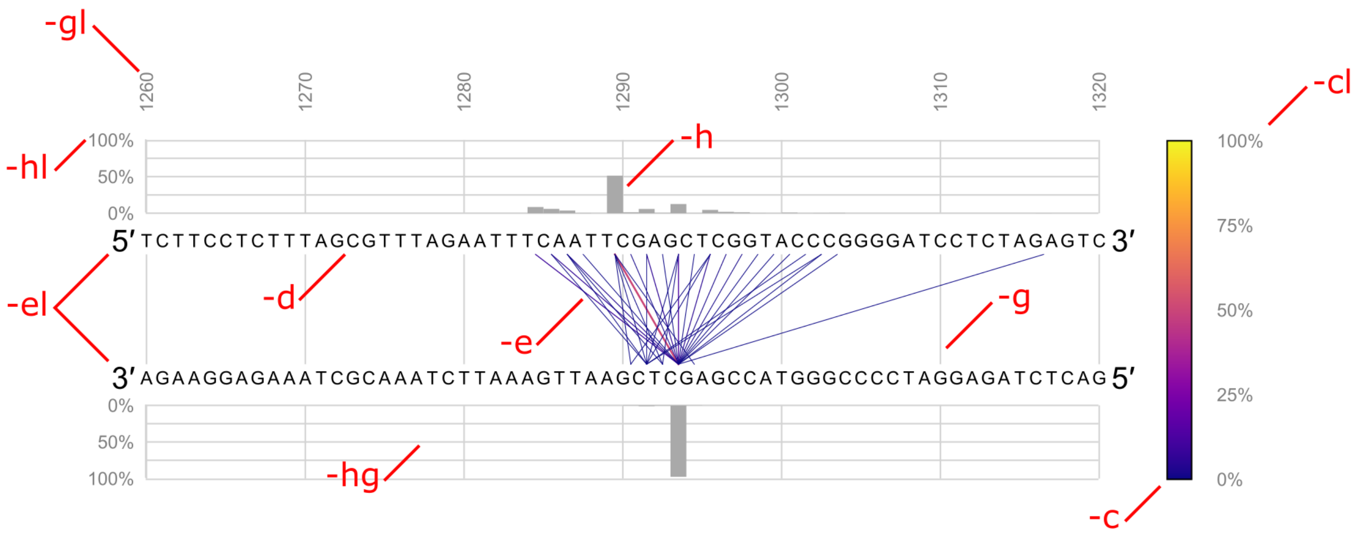


- As shown in the figure above, optional arguments are grouped by the plot feature they act upon. For example, `-gl_i` controls the grid label interval. The following table shows all the optional arguments by feature group:

| Root argument | Feature | Instances |
|---|---|---|

| Root argument | Feature | Instances |
| --- | --- | --- |
| `-d`, `--dna` | DNA sequence | `-d_m`, `--dna_mode`<br>`-d_s`, `--dna_size`<br>`-d_c`, `--dna_colour`<br>`-d_rg`, `--dna_rel_gap` |
| `-el`, `--end_label` | End label (i.e. 5′ and 3′) | `-el_v`, `--end_label_vis`<br>`-el_s`, `--end_label_size`<br>`-el_c`, `--end_label_colour`<br>`-el_rg`, `--end_label_rel_gap`<br>`-el_p`, `--end_label_position` |
| `-g`, `--grid` | Grid (sequence position) | `-g_v`, `--grid_vis`<br>`-g_s`, `--grid_size`<br>`-g_c`, `--grid_colour`<br>`-g_i`, `--grid_interval` |
| `-gl`, `--grid_label` | Grid label (sequence position) | `-gl_v`, `--grid_label_vis`<br>`-gl_s`, `--grid_label_size`<br>`-gl_c`, `--grid_label_colour`<br>`-gl_i`, `--grid_label_interval`<br>`-gl_rg`, `--grid_label_rel_gap` |
| `-c`, `--cbar` | Colourbar | `-c_v`, `--cbar_vis`<br>`-c_rp`, `--cbar_rel_pos`<br>`-c_s`, `--cbar_size` |
| `-cl`, `--cbar` | Colourbar label | `-cl_v`, `--cbar_label_vis`<br>`-cl_s`, `--cbar_label_size`<br>`-cl_c`, `--cbar_label_colour`<br>`-cl_i`, `--cbar_label_interval`<br>`-cl_rg`, `--cbar_label_rel_gap` |
| `-e`, `--event` | Event (linkage lines) | `-e_mis`, `--event_min_size`<br>`-e_mas`, `--event_max_size`<br>`-e_c`, `--event_colourmap`<br>`-e_r`, `--event_range`<br>`-e_orv`, `--event_outside_range_vis`<br>`-e_o`, `--event_opacity`<br>`-e_so`, `--event_stack_order` |
| `-h`, `--hist` | Histogram | `-h_v`, `--hist_vis`<br>`-h_r`, `--hist_range`<br>`-h_bw`, `--hist_bin_width`<br>`-h_c`, `--hist_colour`<br>`-h_rh`, `--hist_rel_height`<br>`-h_rg`, `--hist_rel_gap`<br>`-h_pbg`, `--hist_pc_bar_gap`<br>`-h_o`, `--hist_overhang` |

| Root argument | Feature | Instances |
|---|---|---|
| `-hl`, `--hist_label` | Histogram label | `-hl_v`, `--hist_label_vis`<br>`-hl_s`, `--hist_label_size`<br>`-hl_c`, `--hist_label_colour`<br>`-hl_i`, `--hist_label_interval`<br>`-hl_rg`, `--hist_label_rel_gap`<br>`-hl_p`, `--hist_label_position`<br>`-hl_zv`, `--hist_label_zero_vis` |
| `-hg`, `--hist_grid` | Histogram grid | `-hg_v`, `--hist_grid_vis`<br>`-hg_s`, `--hist_grid_size`<br>`-hg_c`, `--hist_grid_colour`<br>`-hg_i`, `--hist_grid_interval` |

- Many optional arguments share the same form, the most common of these are listed below (for a full list with descriptions use the `-h` or `--help` flag):

| Argument ending | Description | Accepted values |
|---|---|---|
| `v`, `_vis` | Controls whether the feature should be displayed | 'show', 'hide' |
| `s`, `size` | Line widths (in pixel units) for lines or font sizes for text | Non-negative integers |
| `c`, `colour` | Colour of the feature | Colour names (e.g. "black"), hex values (e.g. "#16C3D6") or RGB values in the range 0-255 (e.g. "rgb(128,0,128)") |
| `i`, `interval` | Spacing between numeric features (e.g. grid lines) | Non-negative integers |
| `rg`, `rel_gap` | Gap between the feature and the main strand linkage plot. Specified as a proportion of the width or height of the image. | Floating-point value in the range 0-1 |

### Generating heatmap plots (CSV)

#### Basic use

- Heatmaps can be exported to CSV directly from CSI summary and individual results files using heatmapcsv.py.
- At a minimum, heatmapcsv.py requires arguments specifying the path to a CSI summary or individual results file (`-d` or `--data_file` argument) and the output CSV path (`-o` or `--out_path` argument).
- For example, the following command will generate a heatmap file using default parameters:

```
python .\src\heatmapcsv.py -d .\data\ex_consensus_summary.csv -o
.\data\output_heatmap.csv
```

- Each column of the output heatmap corresponds to a top-strand position and similarly, each row corresponds to a bottom-strand position.
- With default parameters, the final row and column in the heatmap correspond to the sum of all events at that position.
- The total number of events in the heatmap is recorded below the heatmap in the first column.

##### Advanced control

- Further control over the output CSV format can be achieved using the optional arguments listed below:

| Argument | Description | Default value |
|---|---|---|
| `-r`, `--ref_path` | Path to reference sequence file | NA |
| `-ad`, `--append_datetime` | Append time and date to all output filenames (prevents accidental file overwriting) | NA |
| `-pr`, `--pos_range` | Minimum and maximum top and bottom strand positions within the reference sequence to display. Specified as four integer numbers in the order minimum_top maximum_top minimum_bottom maximum_bottom (e.g. -pr 100 200 400 500). If unspecified, the full reference range will be used | 0 0 0 0 |
| `-eldp`, `--event_label_decimal_places` | Number of decimal places to use when displaying event frequencies | 1 |
| `-sv`, `--sum_vis` | Controls whether the sum row and columns are displayed. Must be either "show" or "hide" (e.g. -sv "show") | "show" |
| `-cv`, `--count_vis` | Controls whether the total number of events is displayed underneath the map. Must be either "show" or "hide" (e.g. -cv "show") | "show" |

#### Generating heatmap plots (SVG)

##### Basic use

- Heatmaps can be exported to SVG directly from CSI summary and individual results files using heatmapsvg.py.
- At a minimum, heatmapsvg.py requires arguments specifying the path to a CSI summary or individual results file (`-d` or `--data_file` argument) and the output CSV path (`-o` or `--out_path` argument).
- Without a position range specified (`-pr` or `--pos_range`) the plot will be generated for the top and bottom strand ranges covering all identified cleavage events. The aspect ratio of each event cell is always square.

- For example, the following command will generate a heatmap figure using default parameters:

```
python .\src\heatmapsvg.py -d .\data\ex_consensus_summary.csv -o
.\data\output_heatmap.svg
```

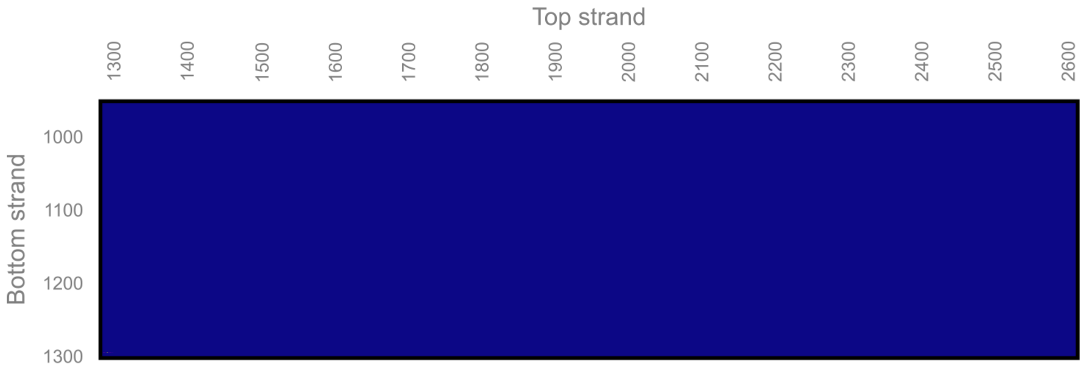


- Note: In this example, the identified events span a large region; however, the vast majority of events are confined to a small position range, so are difficult to see. To zoom in on a region, we can use the optional arguments (see "Advanced control" below).

#### Advanced control

- As with strand linkage plots, comprehensive control over the output heatmaps can be achieved using optional command line arguments.
- Full descriptions for these arguments can be viewed using the `-h` or `--help` flag.
- The following figure uses optional arguments to zoom in on a specific region of the heatmap (`-pr 1280 1300 1285 1300`), renders the grid (`-g_v show`), reduces the grid label interval (`-gl_i 5`) and renders the percentage of events corresponding to each cell (`-el_v show`) and the sum at each position (`-s_v show`):

```
python .\src\heatmapsvg.py -d .\data\ex_consensus_summary.csv -o
.\data\output_heatmap.svg -pr 1280 1300 1285 1300 -g_v show -gl_i 5 -el_v show -
s_v show
```

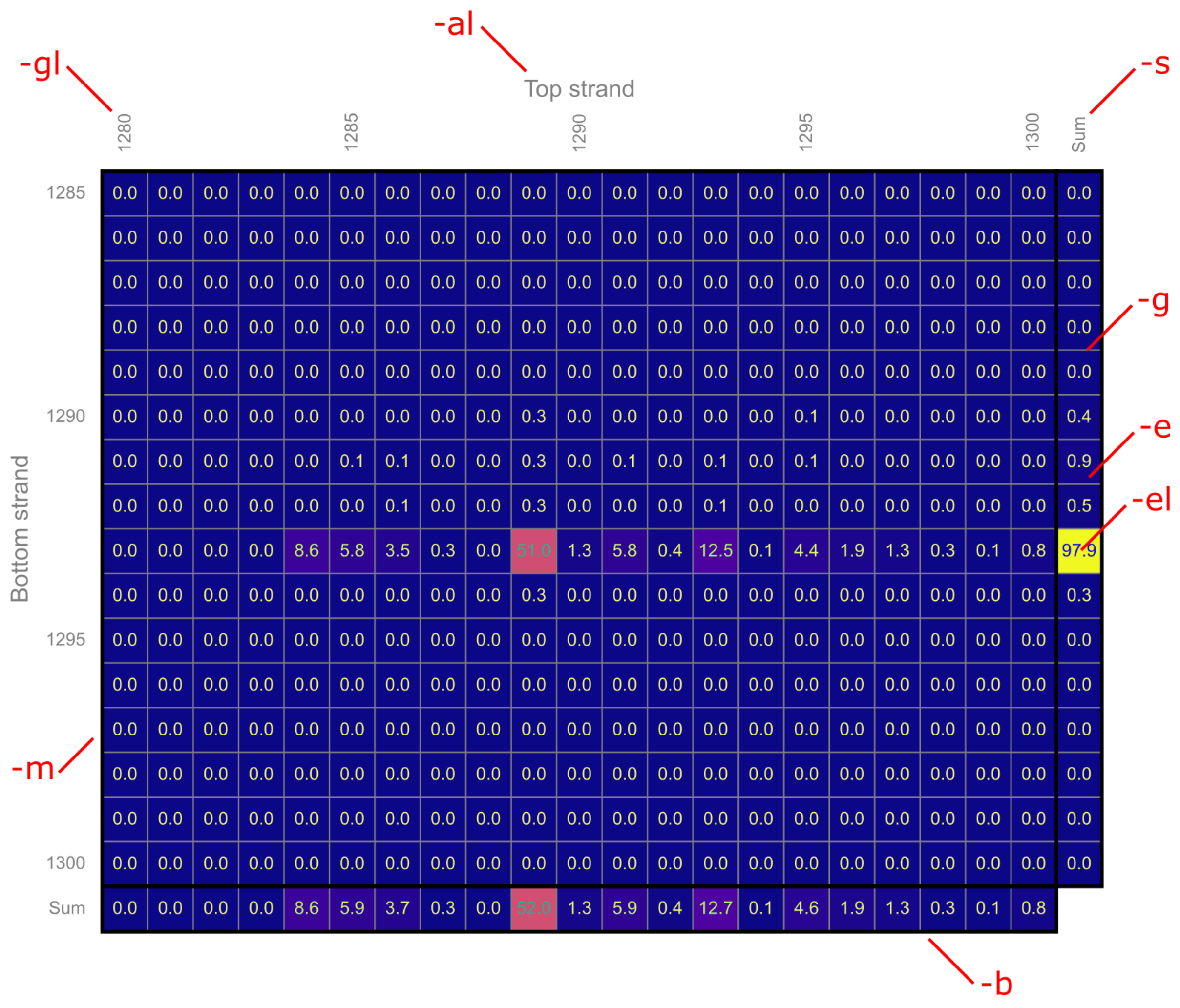


- As shown in the figure above, optional arguments are grouped by the plot feature they act upon. For example, `-gl_i` controls the grid label interval. The following table shows all the optional arguments by feature group:

| Root argument | Feature | Instances |
|---|---|---|
| `-m`, `--map` | Map | `-m_rp`, `--map_rel_pos` |
| `-b`, `--border` | Border | `-b_v`, `--border_vis`<br>`-b_s`, `--border_size`<br>`-b_c`, `--border_colour` |
| `-al`, `--axis_label` | Axis label | `-al_v`, `--axis_label_vis`<br>`-al_s`, `--axis_label_size`<br>`-al_c`, `--axis_label_colour`<br>`-al_g`, `--axis_label_gap` |
| `-g`, `--grid` | Grid | `-g_v`, `--grid_vis`<br>`-g_s`, `--grid_size`<br>`-g_c`, `--grid_colour`<br>`-g_i`, `--grid_interval` |

| Root argument | Feature | Instances |
|---|---|---|
| `-gl`, `--grid_label` | Grid label | `-gl_v`, `--grid_label_vis`<br>`-gl_s`, `--grid_label_size`<br>`-gl_c`, `--grid_label_colour`<br>`-gl_i`, `--grid_label_interval`<br>`-gl_g`, `--grid_label_gap` |
| `-e`, `--event` | Event | `-e_c`, `--event_colourmap` |
| `-el`, `--event_label` | Event label | `-el_v`, `--event_label_vis`<br>`-el_s`, `--event_label_size`<br>`-el_c`, `--event_label_colour`<br>`-el_dp`, `--event_label_decimal_places`<br>`-el_zv`, `--event_label_zeros_vis` |
| `-s`, `--sum` | Sum | `-s_v`, `--sum_vis` |

- Argument endings (e.g. `vis` and `interval`) are similar to those listed for strand linkage plots.

### Output file formats

#### CSI summary file

- Summary CSV files contain a pair of information rows (second row containing just bottom-strand sequence) for each unique restriction site identified in the consensus sequence(s).
- The final row of each summary file reports the number of consensus sequences for which cleavage events could not be determined.
- An example summary file is included in the "data" folder ("ex_consensus_summary.csv").
- Summary files include the following columns:

| Column | Description |
|---|---|
| "TYPE" | Type of cleavage event (either "Blunt end", "3′ overhang" or "5′ overhang"). |
| "COUNT" | Number of identified events matching this cleavage event (% of total identified events shown in parenthesis). |
| "EVENT_%" | Percentage of all identified events (i.e. doesn't include unmatched sequences) corresponding to this event. |
| "TOP_POS" | Position of the top-strand cleavage event. |
| "BOTTOM_POS" | Position of the bottom-strand cleavage event. |
| "SPLIT_SEQ" | `TRUE` if the cleavage event spanned the start/end of the reference sequence, `FALSE` otherwise. |

| Column | Description |
| --- | --- |
| "TOP_LOCAL_SEQ" | Sequence immediately 5′ and 3′ of the cleavage event on the top strand. The number of nucleotides included either side is determined by the `-lr` (or `--local_r`) command line argument. |
| "BOTTOM_LOCAL_SEQ" | Sequence immediately 5′ and 3′ of the cleavage event on the bottom strand. The number of nucleotides included either side is determined by the `-lr` (or `--local_r`) command line argument. |
| "SEQUENCE" | Complete top and bottom strand sequences spanning both cleavage sites. The first row corresponds to the top strand and the second to the bottom strand. Cleavage sites on each strand are represented by the "\|" character. |

#### CSI individual results file

- Individual results files contain a pair of rows (second row containing just bottom-strand sequence) for each consensus sequence processed.
- An example individual results file is included in the "data" folder ("ex_consensus_individual.csv").
- individual results files include the following columns:

| Column | Description |
| --- | --- |
| "INDEX" | Index of this sequence in the input consensus sequence file. Numbering starts at 1. |
| "HEADER" | Header text for this sequence. This is any text on the ">" line imediately preceeding the sequence in the FASTA file. |
| "TYPE" | Type of cleavage event (either "Blunt end", "3′ overhang" or "5′ overhang"). |
| "TOP_LOCAL_SEQ" | Position of the top-strand cleavage event. |
| "BOTTOM_POS" | Position of the bottom-strand cleavage event. |
| "SPLIT_SEQ" | `TRUE` if the cleavage event spanned the start/end of the reference sequence, `FALSE` otherwise. |
| "TOP_LOCAL_SEQ" | Sequence immediately 5′ and 3′ of the cleavage event on the top strand. The number of nucleotides included either side is determined by the `-lr` (or `--local_r`) command line argument. |
| "BOTTOM_LOCAL_SEQ" | Sequence immediately 5′ and 3′ of the cleavage event on the bottom strand. The number of nucleotides included either side is determined by the `-lr` (or `--local_r`) command line argument. |

| Column | Description |
| --- | --- |
| "SEQUENCE" | Complete top and bottom strand sequences spanning both cleavage sites. The first row corresponds to the top strand and the second to the bottom strand. Cleavage sites on each strand are represented by the "\|" character. |
