## Supplementary material for "ENDO-Pore: High-throughput linked-end mapping of single DNA cleavage events using nanopore sequencing": Wet lab protocol

### ENDO-Pore LAB PROTOCOL

#### DNA Cleavage Mapping by Nanopore Sequencing

##### End Repair

Reagents:

- NEBNext® Ultra™ II End Repair/dA-Tailing Module (E7546)

Protocol

1. In a 0.5 mL tube, mix in the following order (place on ice after addition of Enzyme mix):

| | |
|---|---|
| DNA (1 µg) | 50 µl |
| NEBNext Ultra II End Prep Reaction Buffer | 7 µl |
| NEBNext Ultra II End Prep Enzyme Mix | 3 µl |
| **Total** | **60 µl** |

2. Incubate 30 minutes at 20 °C followed by 30 minutes at 65 °C (Use a thermal cycler, heated lid at 75 °C).
3. Transfer the tubes to ice and stop reaction immediately by purification with the "DNA Clean & Concentrator-5 kit" from. Elute in 15 µl of Milli-Q water.

##### Ligation

Reagents:

- T4 DNA ligase (NEB-M0202S)

Protocol:

1. In a 1.5 mL Eppendorf DNA LoBind tube, mix in the following order:

   Perform ligation with 100 ng of linearized plasmid vector and an insert-to-vector ratio of 3:1. The values below correspond to the chloramphenicol resistance cassette (Cm) and a plasmid of 2.6 kb.

| | |
|---|---|
| Milli-Q water | x µl |
| T4 DNA Ligase Buffer (10x) | 2 µl |
| Plasmid DNA (100 ng) | x µl |
| e.g. Cm cassette (86 ng) | y µl |
| T4 DNA ligase (400 units) | 1 µl |
| **Total** | **20 µl** |

2. Incubate at 16 °C for 18 hours followed by heat inactivation 10 minutes at 65 °C.

**Transformation**

Reagents:

- One Shot® OmniMAX™ 2-T1R chemically competent cells (ThermoFisher)
- SOC Media
- LB agar plates supplemented with chloramphenicol (34 µg/ml)

Protocol

1. Thaw, on ice, one vial of One Shot® OmniMAX™ 2-T1R chemically competent cells for each transformation.
2. Add 5 µl of the ligation into a vial of One Shot® cells and mix gently. Do not mix by pipetting up and down.
3. Incubate the vial(s) on ice for 30 minutes.
4. Heat-shock the cells for 30 seconds at 42 °C without shaking.
5. Remove the vial(s) from the 42 °C bath and place them on ice for 2 minutes.
6. Add 250 µl of pre-warmed S.O.C. Medium to each vial.
7. Cap the vial(s) tightly and shake horizontally at 37 °C for 1 hour at 225 rpm in a shaking incubator.
8. Plate the culture on a LB agar plate supplemented with chloramphenicol. Use 90 µl of culture per plate for a total of 3 plates. Additionally, perform a serial dilution (5 µl of culture in 45 µl of LB, two consecutive dilutions). Plate the second dilution culture on a LB agar plate supplemented with chloramphenicol. Perform the dilutions in duplicate.
9. Invert the plate(s) and incubate at 37 °C overnight.
10. Count the colonies present on the two dilution plates. Multiply the number of colonies observed by 540 (27x10) to have an estimate of the CFU per transformation.

**Plate scrapping/Midiprep**

Reagents:

- HiSpeed Midiprep Kit (QIAGEN-13643)

Protocol:

1. Add 2 mL of liquid LB to each plate. Scrape the colonies and transfer to a 15 mL falcon tube.
2. Centrifuge cells for 15 minutes at 5000 rpm at 4 °C. Discard the supernatant.
3. Purify the plasmid from the cell pellet using the HiSpeed Midiprep Kit. Follow the instructions from the kit. In the last step elute in 0.75-1 mL of Milli-Q water.
4. Measure DNA concentration using a Qubit 4 Fluorometer (Thermo Fisher Scientific).

**Rolling Circle Amplification (RCA)**

Reagents:

- EquiPhi29 DNA Polymerase (Thermo-A39391)
- 10 mM dNTPs Mix (Thermo-R0191)
- Exo-Resistant random primer (Thermo-SO181)

Protocol

1. In a 0.5 mL tube, mix in the following order (DNA mix):

| | |
|---|---|
| Nuclease Free water | 5 µl |
| EquiPhi Reaction buffer | 1 µl |
| Exo-resistant random primer | 2 µl |
| Plasmid DNA (5 ng/ µl) | 2 µl |
| **Total** | **20 µl** |

2. Incubate for 3 minutes at 95 °C in a thermal cycler. Transfer immediately to ice and incubate for 3 to 5 minutes.
3. In a separate 0.5 mL tube, mix in the following order:

| | |
|---|---|
| Nuclease Free water | 20.6 µl |
| EquiPhi Reaction buffer | 3 µl |
| DTT (100 mM) | 0.4 µl |
| dNTPs Mix (10 mM) | 4 µl |
| DNA mix (from step 1,2) | 10 µl |
| EquiPhi29 DNA pol | 2 µl |
| **Total** | **40 µl** |

4. Incubate for 2 hours at 45 °C. Inactivate EquiPhi29 DNA pol for 10 minutes at 65 °C.
5. Transfer the reaction to a 1.5 mL Eppendorf DNA LoBind tube.
6. Purify DNA by using 40 µl of AMPure beads Mix by flicking the tube.
7. Incubate for 5 minutes in a rotator at room temperature (Prepare 700 µl of 70% (v/v) ethanol per sample).
8. Spin down the sample and place in magnetic rack. Wait until beads are fully captured and remove the supernatant.
9. Wash the with 200 µl of 70% (v/v) ethanol. Leave the tubes in the magnetic rack and avoid disturbing the pellet. Remove the supernatant (use a new tip per sample).
10. Repeat the previous step.

11. Spin down, place in magnetic rack and remove residual ethanol. Leave the beads to dry for 10 seconds (Do not exceed drying time, this can lead to a loss of DNA)
12. Add 200 µl of nuclease water. Resuspend the beads by flicking the tube gently. Leave at room temperature for 10 minutes. Mix constantly by gently flicking the tube.
13. Place the tube in the magnetic rack. Transfer the supernatant to a new 1.5 mL Eppendorf DNA LoBind tube. Pipette slowly to avoid disturbing the beads. It is OK to leave some supernatant in the tube to avoid taking beads into the purified DNA.
14. Measure DNA concentration using a Qubit 4 Fluorometer (Thermo Fisher Scientific).

**Debranching using T7 endonuclease**

Reagents:

T7 Endonuclease I (NEB-M0302)

Proteinase K (NEB- P8107S)

Protocol:

1. In a 1.5 mL Eppendorf DNA LoBind tube, mix in the following order:

| | |
|---|---|
| DNA (7 µg) | 128 µl |
| NEB buffer 2 | 15 µl |
| **Total** | **143 µl** |

2. Mix by flicking the tube.
3. Start the reaction by adding 7 µl of T7 endonuclease I (use 1 µl of enzyme per µg of DNA). Incubate at 37 °C for 15 minutes. If processing multiple samples, start the reaction at different points so the reaction time is equal for all samples.
4. Stop reaction by adding 1 µl of proteinase K (0.8 units) and incubate for 5 minutes at 37 °C.
5. Purify DNA by using 75 µl of AMPure beads (If the volume of the T7 Endonuclease reaction was changed, adjust the amount of AMPure beads to 0.5 volumes of the reaction). Mix by flicking the tube.
6. Incubate for 5 minutes in a rotator at room temperature (Prepare 500 µl of 70% (v/v) ethanol per sample).
7. Spin down the sample and place in magnetic rack. Wait until beads are fully captured and remove the supernatant.
8. Wash the with 300 µl of 70% (v/v) ethanol. Leave the tubes in the magnetic rack and avoid disturbing the pellet. Remove the supernatant (use a new tip per sample).
9. Repeat the previous step.

10. Spin down, place in magnetic rack and remove residual ethanol. Leave the beads to dry for 10 seconds (Do not exceed drying time, this can lead to a loss of DNA).
11. Add 67 µl of nuclease water.  Resuspend the beads by flicking the tube gently. Leave at room temperature for 2 minutes.
12. Place the tube in the magnetic rack. Transfer the supernatant to a new 1.5 mL Eppendorf DNA LoBind tube. Pipette slowly to avoid disturbing the beads.
13. Measure DNA concentration using a Qubit 4 Fluorometer (Thermo Fisher Scientific).

**<u>Short Read Eliminator XS</u>**

Reagent:

Short Read Eliminator XS (Circulomics- SKU SS-100-121-01)

Protocol:

1. Adjust the DNA sample to a total volume of 60 µL and a Qubit DNA concentration of between 25 – 150 ng/µL.
2. Pipette sample into a 1.5 mL Eppendorf DNA LoBind tube.
3. Add 60 µL of Buffer SRE XS to the sample. Mix thoroughly by gently tapping the tube.
4. Load tube into centrifuge with the hinge facing toward the outside of the rotor.
5. Centrifuge at 10,000 x g for 30 minutes at room temperature
6. Carefully remove supernatant from tube without disturbing the DNA pellet. The DNA pellet will have formed on the bottom of the tube under the hinge region.
5. Add 200 µL of the 70% EtOH wash solution to tube and centrifuge at 15,000 x g for 4 minutes at room temperature. Do not tap or mix after adding 70% (v/v) ethanol. Place tube directly into centrifuge.
6. Carefully remove wash solution from tube without disturbing the DNA pellet.
7. Repeat step 5 and 6.
8. Add 60 µL of Buffer EB to the tube and incubate at 50 °C for 1 hour. Tap the tube gently every 10-15 minutes to aid pellet resuspension.
11. After incubation, gently tap the tube to ensure that the DNA is properly resuspended and mixed.
12. Measure DNA concentration using a Qubit 4 Fluorometer (Thermo Fisher Scientific). Perform two measurements per sample. The values should be similar. If the values differ significatively, mix the sample by tapping and measure again.

**MinION Library prepration**

Follow the instructions in the Native Barcode Protocol (SQ-LSK109, EXP-NBD104/NBD114), with the following modifications:

- Use 1.1 µg of DNA per sample as a starting point for End repair/ dA tailing.
- Use 600 ng of DNA per sample as a starting point for barcode ligation.
