## Supplementary figures for "ENDO-Pore: High-throughput linked-end mapping of single DNA cleavage events using nanopore sequencing"

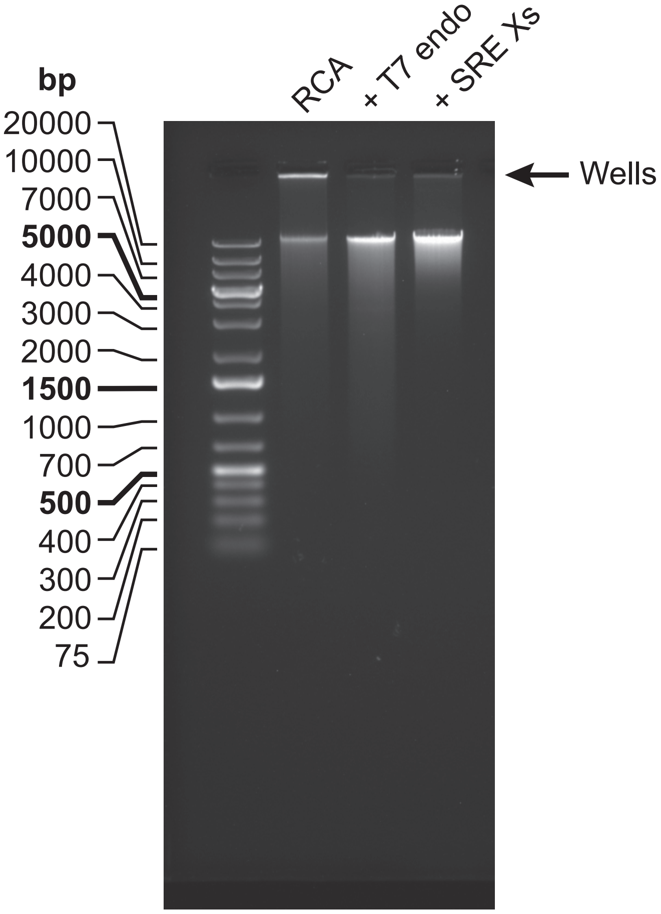


**Supplementary Figure S1. Debranching and DNA size selection following rolling circle amplification.** The branched DNA that results from RCA with random primers runs as a smear on a 0.8% (w/v) agarose gel with much of the DNA stuck in the well. Following debranching by treatment with T7 endonuclease, more of the DNA can enter the gel. A Short Read Eliminator XS Kit (Circulomics) is used to remove DNA <5 kb nearly completely and to deplete DNA <10 kb progressively.

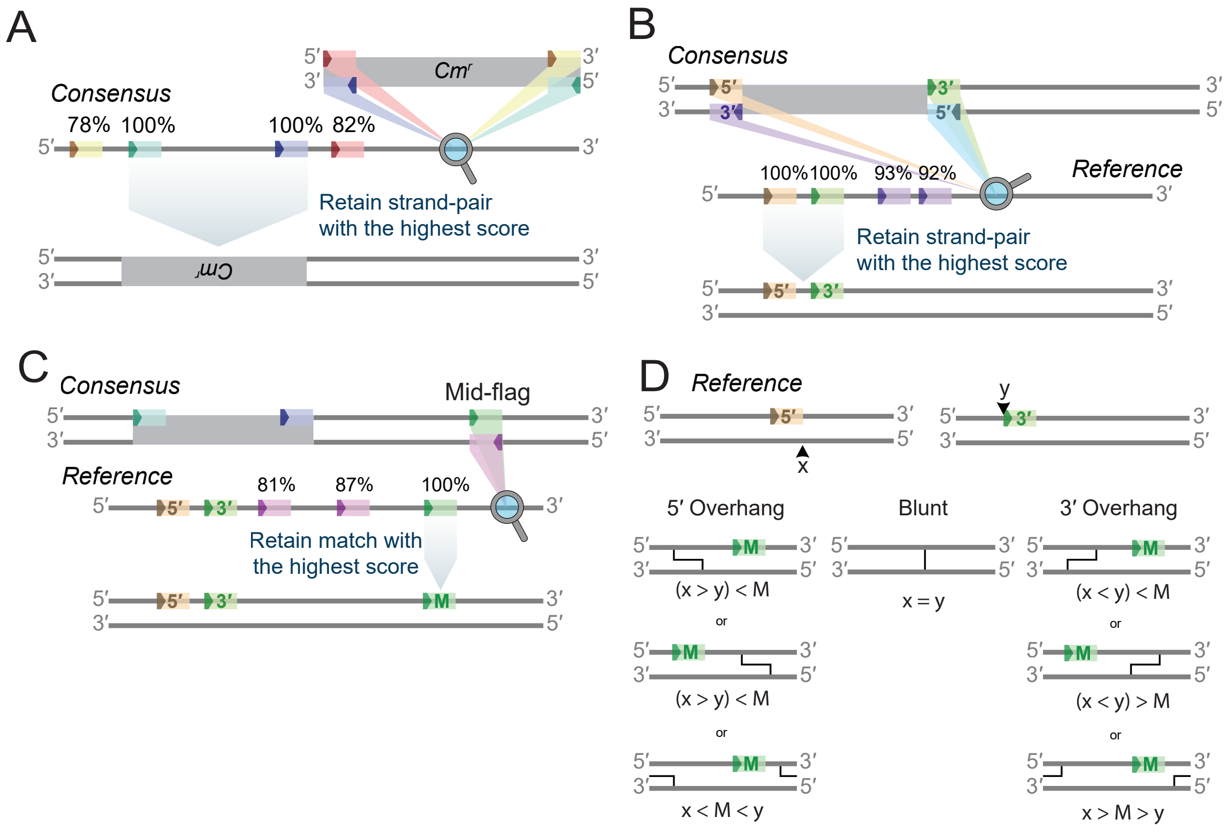


**Supplementary Figure S2. Schematic of the site search logic used by Cleavage Site Investigator.** (**A**) Strand-paired sequences from each end of the chloramphenicol cassette are used to search the consensus sequence generated by C3POa. The pair with the highest sequence identity score to the consensus are retained. (**B**) Strand-paired sequences on each strand of the consensus 5′ and 3′ to the identified chloramphenicol cassette are used to search the reference sequence (e.g. pUC19). The pair with the highest sequence identity score are retained. (**C**) Mid-flag sequences from both strands of the consensus equal in distance from the 5′ and 3′ ends of the chloramphenicol cassette are used to search the reference sequence. The mid-flag sequence with the highest identity score is retained (M). (**D**) The sequences identified in panel B are used to map the cleavage site; the bottom strand cleavage (position x) is mapped from the 3′ terminal end of the 5′ sequence; the top strand cleavage (position y) is mapped from the 5′ terminal end of the 3′ sequence. Based on the relative position of the mid-flag sequence (M), logic rules are then used to identify the type of cleavage generated. Note that possible processing of the DNA ends by the endonuclease need to be considered when interpreting the identity of the observed ends (see Figure 2 and Supplementary Figure S4).

A

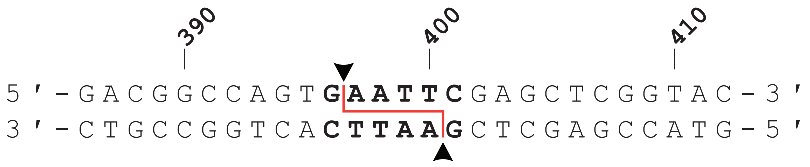


B

```
RESULTS:
    Full sequence frequency:

    TS position: 396
    BS position: 400
    Split seq:   False
    Count:       819/820 (99.9% of events)
    Type:        5' overhang
    Sequence:    5'...CCAGTG|AATT CGAGCT...3'

                 3'...GGTCAC TTAA|GCTCGA...5'


    TS position: 411
    BS position: 400
    Split seq:   False
    Count:       1/820 (0.1% of events)
    Type:        3' overhang
    Sequence:    5'...TGAATT CGAGCTCGGTA|CCCGGG...3'

                 3'...ACTTAA|GCTCGAGCCAT GGGCCC...5'



    Local dinucleotide frequencies:

    AA: 0
    AT: 0            Local 5' nucleotide frequencies:
    AG: 0
    AC: 1            A: 1
    TA: 0            T: 0
    TT: 0            G: 1639
    TG: 0            C: 0
    TC: 0
    GA: 1639
    GT: 0            Local 3' nucleotide frequencies:
    GG: 0
    GC: 0            A: 1639
    CA: 0            T: 0
    CT: 0            G: 0
    CG: 0            C: 1
    CC: 0


    Summary of errors:

    Cassette not found in test:  20
    Cassette ends RC mismatch:   6
    Test not found in reference: 6
    Test ends RC mismatch:       0
    Maximum gap exceeded:        0
    Midpoint not found:          1


    Completed with 33 errors (3.868699%)
```

**Supplementary Figure S3. Cleavage location definition and example output from Cleavage Site Investigator.** (**A**) The EcoRI cleavage site in pUC19. The cleavage positions are defined as 396 on the top strand and 400 on the bottom strand. (**B**) Example output from CSI for a mapping experiment using EcoRI and a repeat cut-off of 7 (N = 820 events). For the cleavage sequences, strand breaks are shown as vertical lines. The low frequency cleavage event represents a DNA cut twice (once at the EcoRI site and once downstream), that then reports as a pseudo-3’ overhang (see Supplementary Figure S4).

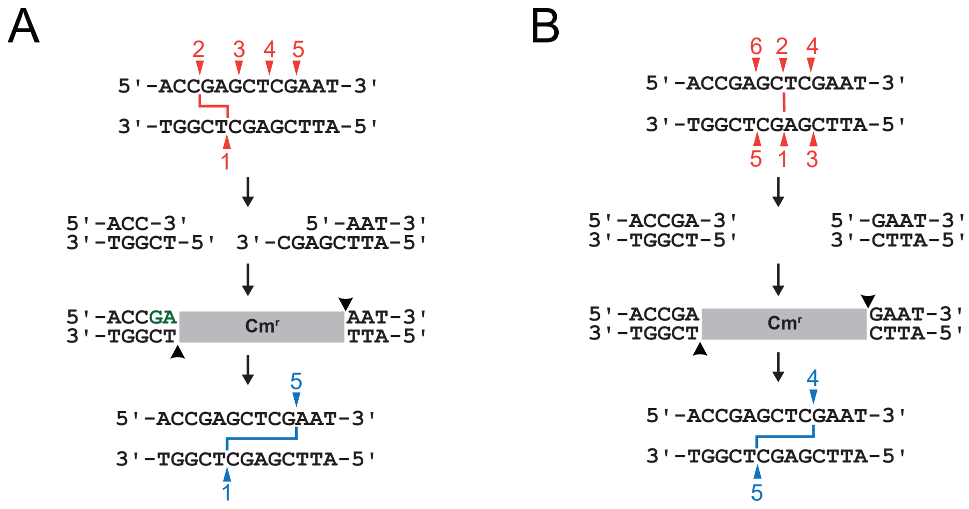


**Supplementary Figure S4. The consequences of DNA end repair are that cleavage sites identified are always those closest to the 3′ end of each strand.** (**A**) Example of 5′ - 3′ end processing. Following initial cleavage (events 1,2) that produces a 2 nt 5′ overhang, further nicking of the top strand (events 3 → 5) produces a 3′ overhang on one end. End repair, cassette ligation and sequencing identify the cleavage sites as events 1 and 5 only (i.e., a 5 nt 3′ overhang). (**B**) Example of an enzyme that moves position and cuts again. Following initial cleavage (events 1,2) that produces a blunt end, the enzyme moves and makes two further blunt end cuts (events 3,4 and 5,6). End repair, cassette ligation and sequencing identify the cleavage sites as events 4 and 5 only (i.e. a 4 nt 3′ overhang). See main text and Figure 2 for further details.

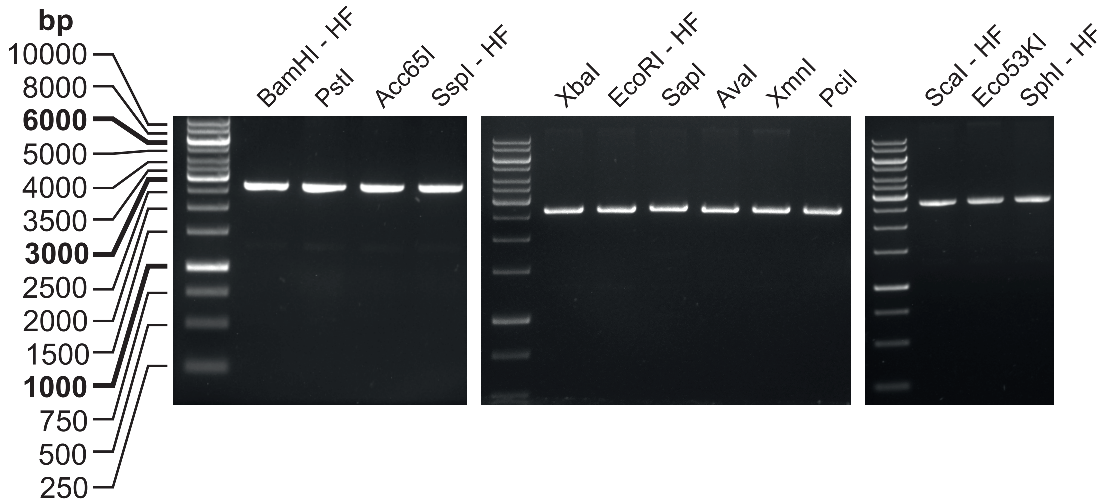


**Supplementary Figure S5. DNA cleavage by commercial Type II restriction endonucleases for ENDO-Pore sequencing.** pUC19 was incubated with the enzymes indicated according to the manufacturer's recommendations (Materials and Methods).

| | CSI sequence | basecalling error | ≥1 | ≥2 | ≥3 | ≥4 | ≥5 | ≥6 | ≥7 | ≥8 | ≥9 |
|---|---|---|---|---|---|---|---|---|---|---|---|
| **XmnI** | 5'..AAA A\|CGTT..3'<br>3'..TTT\|T GCAA..5' | $\Delta$1 bp<br>AAAAAG......CTCGTT<br>TTTTTC......GAGCAA | 9.6%<br>(27542) | 8.1%<br>(18397) | 6.6%<br>(10990) | 5.6%<br>(6581) | 4.5%<br>(4069) | 3.7%<br>(2478) | 3.5%<br>(1505) | 2.8%<br>(936) | 1.8%<br>(603) |
| | 5'..AA AA\|CGTT..3'<br>3'..TT\|TT GCAA..5' | $\Delta$2 bp<br>AAAAAG......CTCGTT<br>TTTTTC......GAGCAA | 0.8%<br>(27542) | 0.4%<br>(18397) | 0.2%<br>(10990) | 0.1%<br>(6581) | 0<br>(4069) | 0<br>(2478) | 0.1%<br>(1505) | 0<br>(936) | 0<br>(603) |
| | 5'..A AAA\|CGTT..3'<br>3'..T\|TTT GCAA..5' | $\Delta$3 bp<br>AAAAAG......CTCGTT<br>TTTTTC......GAGCAA | 0.1%<br>(27542) | 0.1%<br>(18397) | 0.1%<br>(10990) | 0<br>(6581) | 0<br>(4069) | 0<br>(2478) | 0.1%<br>(1505) | 0<br>(936) | 0<br>(603) |

| | CSI sequence | basecalling error | ≥1 | ≥2 | ≥3 | ≥4 | ≥5 | ≥6 | ≥7 | ≥8 | ≥9 |
|---|---|---|---|---|---|---|---|---|---|---|---|
| **Eco53KI** | 5'..GAG CT\|C..3'<br>3'..CTC\|GA G..5' | $\Delta$2 bp<br>GAGAGAG......CTCTCTC<br>CTCTCTC......GAGAGAG | 5.6%<br>(4315) | 4.6%<br>(3296) | 3.2%<br>(2322) | 2.4%<br>(1572) | 2.3%<br>(1098) | 2.1%<br>(752) | 1.8%<br>(495) | 2.0%<br>(357) | n.d. |
| | 5'..G AG\|CTC..3'<br>3'..C\|TC GAG..5' | $\Delta$2 bp<br>GAGAGAG......CTCTCTC<br>CTCTCTC......GAGAGAG | 2.3%<br>(4315) | 1.5%<br>(3296) | 1.0%<br>(2322) | 0.5%<br>(1572) | 0.4<br>(1098) | 0.3<br>(752) | 0.2%<br>(495) | 0<br>(357) | n.d. |
| | 5'..GGTGAGT\|ACTCAAC..3'<br>3'..CCACTCA\|TGAGTTG..5' | None<br>ScaI site<br>indexing error | 0.1%<br>(4315) | 0.1%<br>(3296) | 0.1%<br>(2322) | 0.1%<br>(1572) | 0.2%<br>(1098) | 0.3%<br>(752) | 0.4%<br>(495) | 0.3%<br>(357) | n.d. |

| | CSI sequence | basecalling error | ≥1 | ≥2 | ≥3 | ≥4 | ≥5 | ≥6 | ≥7 | ≥8 | ≥9 |
|---|---|---|---|---|---|---|---|---|---|---|---|
| **XbaI** | 5'..TCT\|AG A..3'<br>3'..AGA TC\|T..5' | $\Delta$2 bp<br>TCTAGAGAG......CTCTCTAGA<br>AGATCTCTC......GAGAGAGCT | 4.5%<br>(25915) | 3.3%<br>(17719) | 2.3%<br>(10055) | 1.6%<br>(5694) | 0.9%<br>(3214) | 0.9%<br>(1802) | 0.4%<br>(1037) | 0.5%<br>(612) | n.d. |
| | 5'..T\|CT AGA..3'<br>3'..G GA\|TCT..5' | $\Delta$2 bp<br>TCTAGAGAG......CTCTCTAGA<br>AGATCTCTC......GAGAGAGCT | 3.0%<br>(25915) | 2.2%<br>(17719) | 1.7%<br>(10055) | 1.1%<br>(5694) | 0.7<br>(3214) | 0.5<br>(1802) | 0.3%<br>(1037) | 0<br>(612) | n.d. |

**Supplementary Figure S6. Incorrect cleavage site identification due to basecalling or indexing errors.**

Numbers of events are shown in the brackets. n.d., not determined.

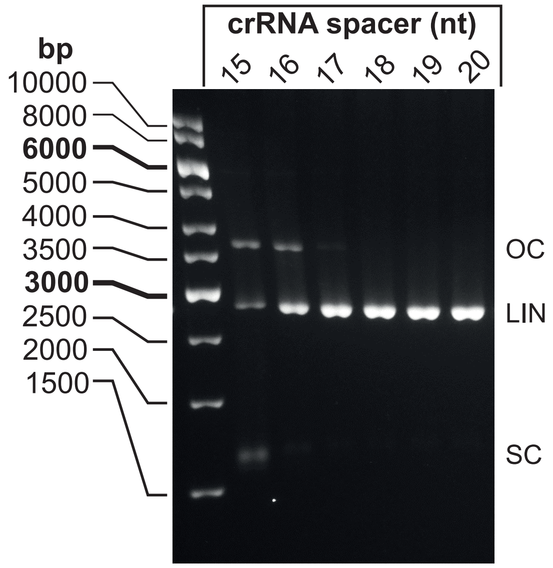


**Supplementary Figure S7. LbCas12a DNA cleavage for ENDO-Pore sequencing.** pSP1 was incubated with CRISPR Cas12a ribonucleoproteins with crRNA with the spacer lengths indicated for 2 hours at 37 °C. Supercoiled (SC) substrate DNA, nicked (open circle, OC) intermediate and linear (LIN) product DNA were separated by agarose gel electrophoresis.

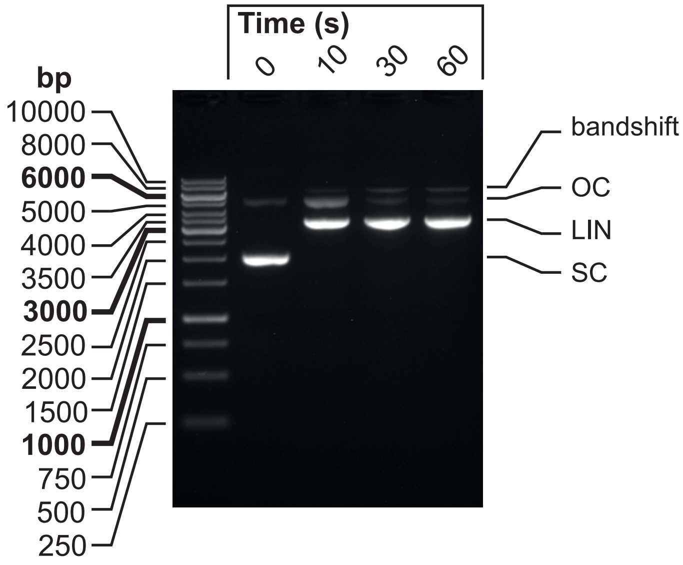


**Supplementary Figure S8. LlaGI DNA cleavage for ENDO-Pore sequencing.** pRMA03S was incubated with LlaGI and 4 mM ATP for the time points indicated at 25 °C. Supercoiled (SC) substrate DNA, nicked (open circle, OC) intermediate and linear (LIN) product DNA were separated by agarose gel electrophoresis. An additional LlaGI-DNA bandshift was observed.

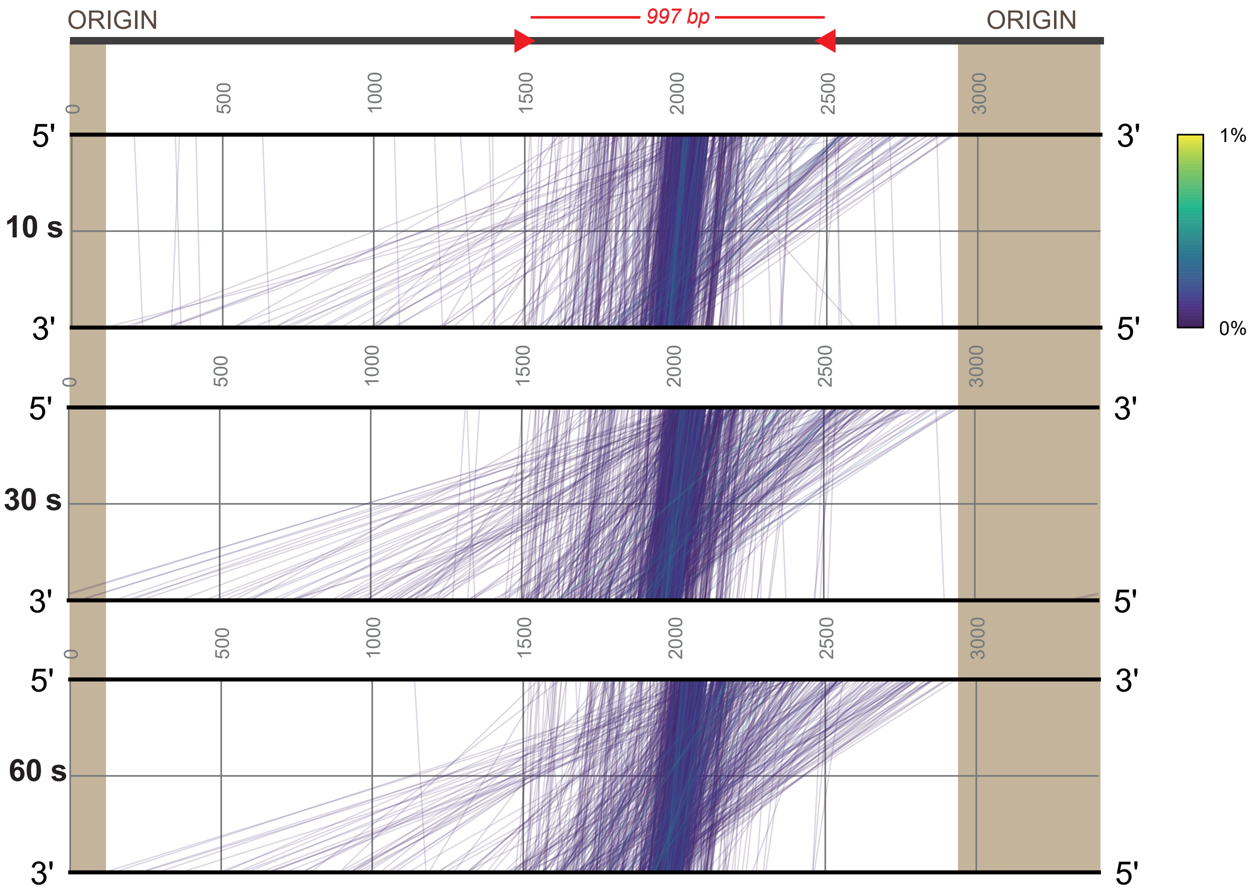


**Supplementary Figure S9. Strand-linkage plots for DNA cleavage by the Type ISP restriction endonuclease LlaGI.** The circular DNA substrate is shown as a black line split arbitrarily within the origin, with the LlaGI sites as red arrowheads indicating the direction of enzyme translocation. The spacing between the sites (red) and the location of the 589 bp Rep origin (brown) are shown. The SLPs for the 10, 30 and 60 s data are shown with frequencies according to the heat map. Note that cleavage sites do not map within the origin since this disrupts plasmid replication and hence transformants are not produced.

| crRNA spacer | RNA sequence (**direct repeat**, **spacer**) |
|---|---|
| **20** | 5′-uAAUUUCUACUAAGUGUAGAUGCGUUUAGAAUUUCAAUUCG-3′ |
| **19** | 5′-uAAUUUCUACUAAGUGUAGAUGCGUUUAGAAUUUCAAUUC-3′ |
| **18** | 5′-uAAUUUCUACUAAGUGUAGAUGCGUUUAGAAUUUCAAUU-3′ |
| **17** | 5′-uAAUUUCUACUAAGUGUAGAUGCGUUUAGAAUUUCAAU-3′ |
| **16** | 5′-uAAUUUCUACUAAGUGUAGAUGCGUUUAGAAUUUCAA-3′ |
| **15** | 5′-uAAUUUCUACUAAGUGUAGAUGCGUUUAGAAUUUCA-3′ |

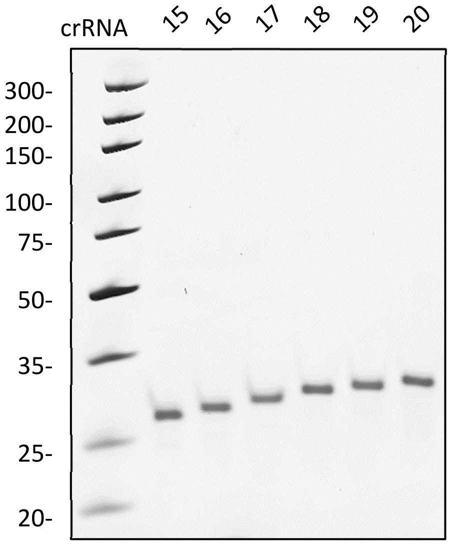


**Supplementary Figure S10. Synthetic LbCas12a crRNA.** The table shows the crRNA sequences. The polyacrylamide gel shows the purified crRNA.
